# Integrating experimental feedback improves generative models for biological sequences

**DOI:** 10.1101/2025.03.31.646327

**Authors:** Francesco Calvanese, Giovanni Peinetti, Polina Pavlinova, Philippe Nghe, Martin Weigt

## Abstract

Generative probabilistic models have shown promise in designing artificial RNA and protein sequences but often suffer from high rates of false positives, where sequences predicted as functional fail experimental validation. To address this critical limitation, we explore the impact of reintegrating experimental feedback into the model design process. We propose a likelihood-based reintegration scheme, which we test through extensive computational experiments on both RNA and protein datasets, as well as through wet-lab experiments on the self-splicing ribozyme from the group I intron RNA family where our approach demonstrates particular efficacy. We show that integrating recent experimental data enhances the model’s capacity of generating functional sequences (e.g. from 6.7% to 63.7% of active designs at 45 mutations). This feedback-driven approach thus provides a significant improvement in the design of biomolecular sequences by directly tackling the false-positive challenge.

## I. INTRODUCTION

Generative probabilistic models for biological sequences, such as proteins and RNA, have recently emerged as promising tools for designing artificial biomolecules [1–4]. These models, particularly family-specific ones like those built using Direct-Coupling Analysis (DCA) [1, 2], as well as more advanced architectures like restricted Boltzmann machines [4], variational autoencoders [2, 5, 6], and protein language models [3], have shown notable success in generating functional sequences. However, a persistent challenge remains: these models often produce a high rate of false positives – sequences generated as potentially functional by the model but failing in experimental tests.

These models are trained on sets of homologous sequences, representing families of sequences with shared evolutionary ancestry. Such families are typically characterized by highly conserved structures and functions, though the sequences themselves may diverge significantly. Multiple-sequence alignments (MSA) [7–10], containing presumably functional sequences from different species, serve as the foundation for training. As a consequence, these models are trained in an unsupervised manner on unlabeled functional sequences, which limits their capacity to differentiate between functional and non-functional variants.

A significant issue in these generative models is the high rate of false positives – sequences deemed functional by the model that fail experimental validation [1, 2]. This limitation arises from the scarce sampling of the viable sequence space in the MSAs, leading to an underrepresentation of functional diversity, and to an intrinsic difficulty in accurately estimating the limitations of functional sequence space.

In this study, we focus on DCA-based Potts models [1, 11, 12] and demonstrate that integrating experimental feedback including false-positive sequences into the training procedure can enhance model accuracy and reduce false-positive rates. By incorporating this feedback through an extension of the maximum-likelihood inference procedure [12], which makes explicit use of the experimental results, we show that false positives from the initial model play a critical role in refining the boundaries of the viable sequence space, thereby improving the model’s performance. An intriguing ingredient to this approach is that the underlying mathematical structure of the model remains unchanged, but the reintegration of experimental data significantly improves parameter learning. This highlights an important insight: the current limitations of generative models stem not necessarily from the limited expressivity of their architectures, but also from the insufficient information content in the original training data. These alignments, representing natural sequences that have diverged through evolution, offer a sparse and incomplete sampling of the functional sequence landscape. Enhancing this landscape with experimental feedback allows for a reliable model, generating a higher fraction of functional sequences (i.e. true positives). Augmenting data quantity and quality at unchanged model complexity turns out to be an efficient strategy.

The paper is organized as follows. In the next *Results* section, we outline the main ideas of the reintegration approach and present extensive tests on diverse RNA and protein families, progressing from purely computational settings to experimental validation : artificial sequences sampled from our models and tested experimentally are the most rigorous possible validation of a generative approach to bio-molecules. We detail the application of our procedure to DCA models in the *Materials and Methods* section, where we also describe the datasets used to evaluate our approach. At the end of the article, we present our main conclusion, and show an outlook of possible extensions of our approach.

## II. RESULTS

Here we propose a method to reintegrate experimental test results into a sequence generative model. Figure 1 illustrates the core idea. In the standard approach, the natural data MSA 𝒟_*N*_ is used to train an initial probabilistic generative model *P* (*<u>a</u>* | *θ*^1^) = *P*^1^(*<u>a</u>*) whose parameters *θ*^1^ are obtained through Maximum Likelihood Estimation (MLE):

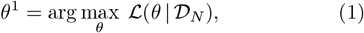

where

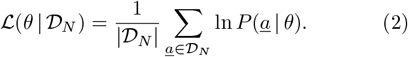

**FIG. 1:**
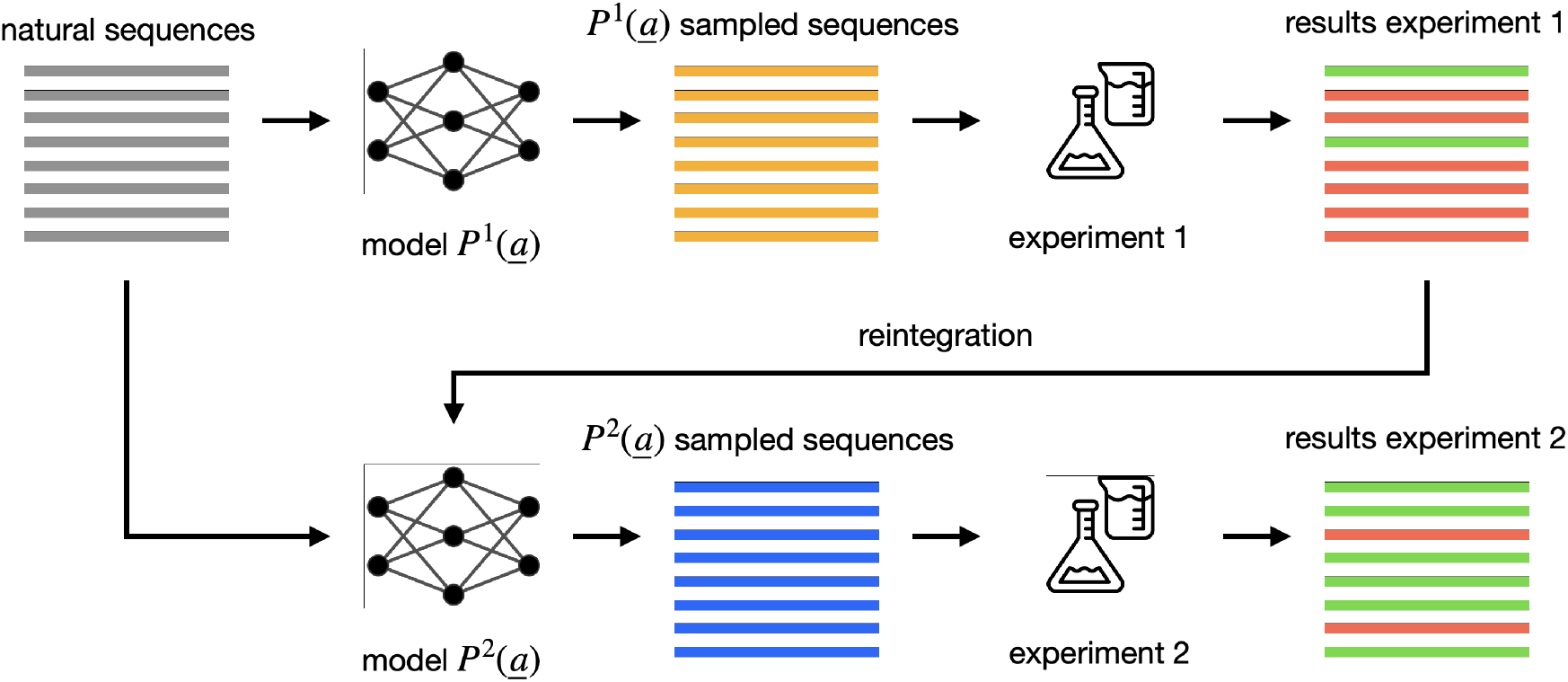
Schematic representation of the reintegration procedure. The original model probability distribution P^1^(<u>a</u>) trained on a natural MSA is used to sample the P^1^-dataset. These sequences are experimentally tested and labeled, and this information is reintegrated into the training of a DCA model P^2^(<u>a</u>). The new P^2^-dataset exhibits enhanced functionality. Note that the mathematical form of P^1^(<u>a</u>) and P^2^(<u>a</u>) is identical, and that the improved generative performance results from refined parameter values learned on enriched data.

Once the model is trained, an set 𝒟_*T*_ of artificial sequences can be sampled from *P*^1^(*<u>a</u>*) and tested experimentally. However, this approach often suffers from a high rate of false-positive sequences in 𝒟_*T*_, i.e. sequences expected to be functional according to *P*^1^(*<u>a</u>*), yet failing experimental tests (indicated in red in Figure 1), cf. [1, 2]. To address this issue, we propose reintegrating the experimental feedback contained in 𝒟_*T*_ into an updated model, *P* (*<u>a</u>* | *θ*^2^) = *P*^2^(*<u>a</u>*). This updated model maintains the same mathematical form and architecture as *P*^1^(*<u>a</u>*) but uses recalibrated parameters inferred leveraging the newly labeled data 𝒟_*T*_. Consequently, *P*^2^(*<u>a</u>*) is expected to generate a higher proportion of functional (true-positive) sequences. To implement this, we update the model parameters optimizing a new objective function:

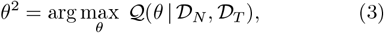

where

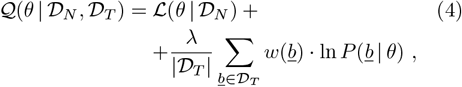

The first contribution to 𝒬 equals the standard loglikelihood for the natural data given in Eq. (2). Maxmizing 𝒬 reduces thus to standard MLE if no experimental data is available, i.e., for 𝒟_*T*_ = ∅. The second contribution is the reintegration term, which acts on the probabilities of the tested sequences in dependence on the adjustment weight *w*, assigned to every sequence in the tested dataset *<u>b</u>* ∈ 𝒟_*T*_ in function of the experimental test result. We require *w*(*<u>b</u>*) to adhere to the following rules:

- Negative Adjustment: *w*(*<u>b</u>*) *<* 0 for sequences failing the experimental functionality test, such that their probability *P*^2^(*<u>b</u>*) is reduced when maximizing 𝒬.
- Positive Adjustment: *w*(*<u>b</u>*) *>* 0 for sequences passing the experimental functionality test, such that their probability *P*^2^(*<u>b</u>*) is increased when maximizing 𝒬.

The specific values of the weights for individual sequences depend on the specific experimental setting, in the easiest case they can be taken to be all of equal absolute value (see below for some more complicated construction removing at least partially biases in the experimental data). The overall intensity of reintegration is controlled by the hyperparameter *λ*; the higher is *λ*, the greater the relative importance assigned to the experimentally labelled dataset 𝒟_*T*_ compared to the natural sequences 𝒟_*N*_. When *λ* = 0, the classical MLE is recovered. If we consider only the functional sequences with *w*(*<u>b</u>*) *>* 0 in 𝒟_*T*_, this procedure is similar to adding them to 𝒟_*N*_ with a *λ*-dependent weight, and performing the standard MLE inference.

The essential difference arises from the inclusion of non-functional sequences with *w*(*<u>b</u>*) *<* 0 in 𝒟_*T*_, which indicate regions of the sequence space that our model should avoid, cf. Fig. 2. Ideally, 𝒟_*T*_ would consist of *P*^1^(*<u>a</u>*) generated sequences, cf. Fig. 1. Consider a sequence *<u>b</u>* that has been generated by *P*^1^(*<u>a</u>*): the sole fact that it was sampled implies that it was assigned a high probability by the *P*^1^(*<u>a</u>*) model. If this sequence fails the experimental test, it is assigned a negative *w*(*<u>b</u>*) *<* 0, and the reintegrated *P*^2^(*<u>a</u>*) model will subsequently assign it a lower probability. This procedure enables the model to correct itself based on the experimental feedback, and to better infer the limits of functional sequence space.

**FIG. 2:**
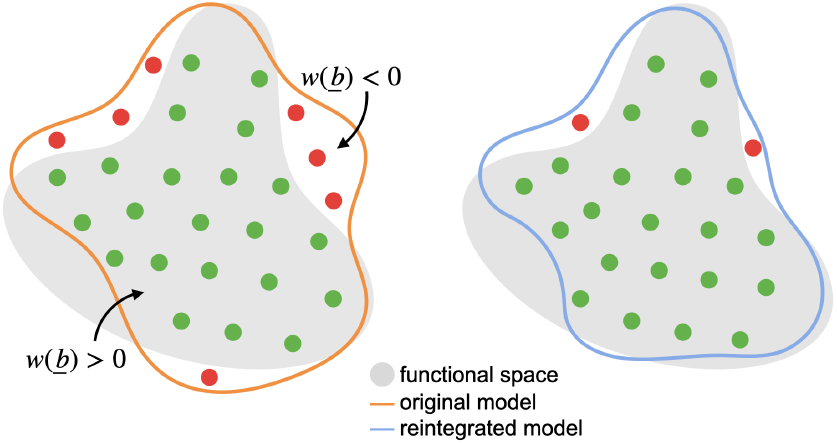
Stylized representation of the effect of the reintegration procedure. Sequences generated by the initial model P^1^ (region inside orange line) that fail experimental tests (region outside grey area) are assigned negative adjustment weights (w < 0), while those that are functional are assigned positive weights (w > 0). The reintegrated model P^2^ is then trained using these adjusted weights, resulting in generated sequences that avoid regions associated with non-functional sequences and concentrate in regions associated with functional sequences (region inside blue line), thereby reducing the fraction of false positives among the generated sequences.

The reintegration set 𝒟_*T*_ is, however, not limited to be comprised of sequences generated by *P*^1^(*<u>a</u>*) but can, in principle, be any functionally labelled dataset. It is, however, intuitively important that negative sequences are close to the functional sequence space. Random sequences, e.g., are almost surely non-functional, but they typically have already very low values of *P*^1^(*<u>a</u>*), and reintegrating them negatively will not improve the description of the positively functional sequence space.

Our tests of this procedure have been carried out on DCA models, since DCA has proved capable of generating functional proteins and RNAs [1, 2]. Another advantage is that we can naturally implement the new objective function 𝒬 in the DCA framework without altering its classical training procedures. A detailed discussion is provided in the *Materials and Methods* section with all analytical derivations detailed in the *Supplementary Section S1*. Note that, in a more generic machine-learning context, the DCA model can be replaced by other generative model architectures, and MLE by the optimization of any loss function, which is additive in data-point specific losses.

In the following, we present computational tests on RNA and protein data, as well as experimental validation performed on group I intron ribozymes.

### A. The effect of the reintegration strength *λ* in Rfam RNA families

To understand the action of the proposed reintegration method, we need to study its performance systematically in function of the reintegration strength *λ*. For this aim, we use three RNA-family MSAs from the Rfam database (cf. *Materials and Methods*). Before performing resource and time consuming experiments (cf. below), we first assess the role of *lambda* via a fully computational approach. As a proxy for the experimental fitness, we employ the negative free energy −*F* of folding a given sequence onto the family’s consensus secondary structure, computed from the Turner Model [13]) implemented in the `RNAeval`function of the ViennaRNA package [14]. Further details on the data and fitness proxy are provided in the *Materials and Methods*.

For the statistical models, we use our recent time-efficient RNA-tailored Edge Activation DCA (eaDCA) [15]. It provides easy access to the inferred models’ Shannon entropy *S* [16] in function of *λ*, and thereby allows to quantify the potential diversity of the sequences generated by the model.

To this end, for each of the three RNA families, we used the Rfam MSA as 𝒟_*N*_ and trained our initial model *P*^1^(*<u>a</u>*) via eaDCA. From this model, we sampled the *P*^1^(*<u>a</u>*)-dataset comprising 2000 sequences, to be used as 𝒟_*T*_ in the reintegration procedure. We measured the `RNAeval` proxy fitness −*F* [14] of these sequences (cf. *Materials and Methods*), and decided (somewhat arbitrarily) to consider all sequences with above-average −*F* as functional, and below-average −*F* as non-functional - the training objective for *P*^2^(*<u>a</u>*) thus being the generation of highly thermo-stable sequences. We thus define a simple adjustment weight *w*(*<u>b</u>*) for all *<u>b</u>* ∈ 𝒟_*T*_ :

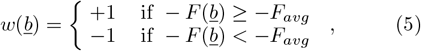

where *F*_*avg*_ is the average `RNAeval`folding free energy evaluated for the 𝒟_*T*_ dataset.

A reasonable range for the reintegration strength *λ* can be chosen using the following consideration: as already mentioned, the case *λ* = 0 is equivalent to training the standard DCA model *P*^1^(*<u>a</u>*) using MLE and the natural MSA 𝒟_*N*_. For *λ* = 1, the two contributions to the objective function 𝒬 in Eq. (4) become equally important, and so do the two datasets 𝒟_*N*_ and 𝒟_*T*_. It is therefore reasonable to explore *λ*s between zero and values slightly larger than one. Note that, for larger values, the learning algorithm starts to have convergence problems (see *Materials and Methods*).

To assess the effect of the reintegration procedure, we monitor the following quantities:

1. *Average Proxy Fitness:* 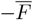 is the average proxy fitness of the sequences in the *P*^2^-dataset. This measures how well our reintegration is pushing the generation towards “functional” sequences according to Eq. (5).
2. *True Positives*: TP is defined as the fraction of sequences in the *P*^2^-dataset that exhibit a fitness score *F* (*<u>b</u>*) ≥ −*F*_*avg*_. In other words, these sequences would have been assigned a positive weight during the reintegration procedure, indicating predicted functionality. This metric quantifies the effectiveness of the approach in reducing false positives.
3. *Model entropy: S* quantifies the diversity of sequences generated by the reintegrated model *P*^2^. A higher entropy suggests a more diverse sequence space.
4. *Average intra-dataset distance:* 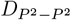 is the average Hamming distance (number of mutations) between pairs of sequences in the *P*^2^-dataset. It quantifies the diversity between generated sequences. This value is to be compared with 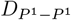, i.e., the average distance in the non reintegrated *P*^1^(*<u>a</u>*)-dataset.
5. *Average minimum distance to functional sequences in* 𝒟_*T*_ : for each sequence in the *P*^2^-dataset, we calculate the Hamming distance to the closest functional sequence in the reintegration dataset 𝒟_*T*_. The average of these is reported as 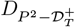. This metric ensures that our model is not merely replicating functional sequences from 𝒟_*T*_.
6. *Average minimum distance to non-functional sequences in* 𝒟_*T*_ : for each sequence in the *P*^2^-dataset, we calculate the Hamming distance to the closest non-functional sequence in the reintegration dataset 𝒟_*T*_. The average of these is reported as 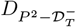. This metric allows us to test if generated sequence avoid the vicinity of non-functional sequences from 𝒟_*T*_, as is to be expected by their negative contribution to the objective 𝒬.

By tracking these quantities, we can evaluate whether the reintegration leads to a better model without overfitting the reintegrated data. The results of these analyses are presented in Table I for the RF00162 RNA family, similar results are observed also for the other families, cf. *Supplementary Section S2* in *Supplementary Tables S1, S2, S3* and *Supplementary Figure S1*.

**TABLE I:** Effect of the reintegration strength for the RF00162 RNA family. Note that the values reported for λ = 0 correspond to the DCA model without reintegration, i.e. −F_avg_ = 14.68 is the threshold value chosen for functional sequences, and the entropy S = 61.92 equals the one of P^1^.

| $\lambda$ | $-\bar{F}$ | TP | $S$ | $D_{P^2-P^2}$ | $D_{P^2-\mathcal{D}_T^+}$ | $D_{P^2-\mathcal{D}_T^-}$ |
| --- | --- | --- | --- | --- | --- | --- |
| 0 | 22.4 | 49.9% | 63.4 | 46.0 | — | — |
| 0.1 | 24.3 | 60.4% | 65.3 | 45.8 | 28.1 | 29.6 |
| 0.5 | 30.2 | 91.3% | 63.2 | 42.2 | 26.7 | 30.5 |
| 1.0 | 34.9 | 99.4% | 53.3 | 39.7 | 25.3 | 31.8 |

In general, we observe that h<u>i</u>gher values of *λ* yield a higher average proxy fitness 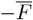, demonstrating that the reintegration procedure effectively enhances the fitness of the generated sequences. For example, for the RF00162 RNA family (Table I), the average proxy fitness increases from 22.4 (*λ* = 0) to 34.9 (*λ* = 1), and the TP fraction improves from 49.9% to 99.4%. However, this gain comes at the expense of model entropy *S* and overall sequence diversity: *S* decreases from 63.4 (*λ* = 0) to 53.3 (*λ* = 1), while the average intra-dataset distance 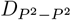 drops from 46.0 (*λ* = 0) to 39.7 (*λ* = 1). Additionally, generated sequences become somewhat closer to the functional sequences in the reintegration dataset 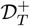, as indicated by a reduction in the average minimum distance 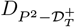 from 28.1 (*λ* = 0.1) to 25.3 (*λ* = 1). The average minimum distance to non-functional sequences 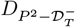 slightly increases from 29.6 (*λ* = 0.1) to 31.8 (*λ* = 1). We were able to perform these tests because we can readily compute the proxy fitness also on the sequences generated from the reintegrated model. In more realistic scenarios, this is not possible without experiments. However, our observations guide the choice of *λ* before doing experiments: it seems reasonable to select a value before a significant loss in diversity occurs or before the training procedure fails to converge.

The effect of reintegration on the proxy fitness 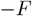 distributions for the three Rfam families is shown in Fig. 3. In all cases, the *P*^2^-dataset samples exhibit a significant shift to higher proxy fitness compared to the *P*^1^-dataset (indicated by *λ* = 0). We find in particular, that “non-functional” sequences with proxy fitnesses below −*F*_*avg*_ become very rare.

**FIG. 3:**
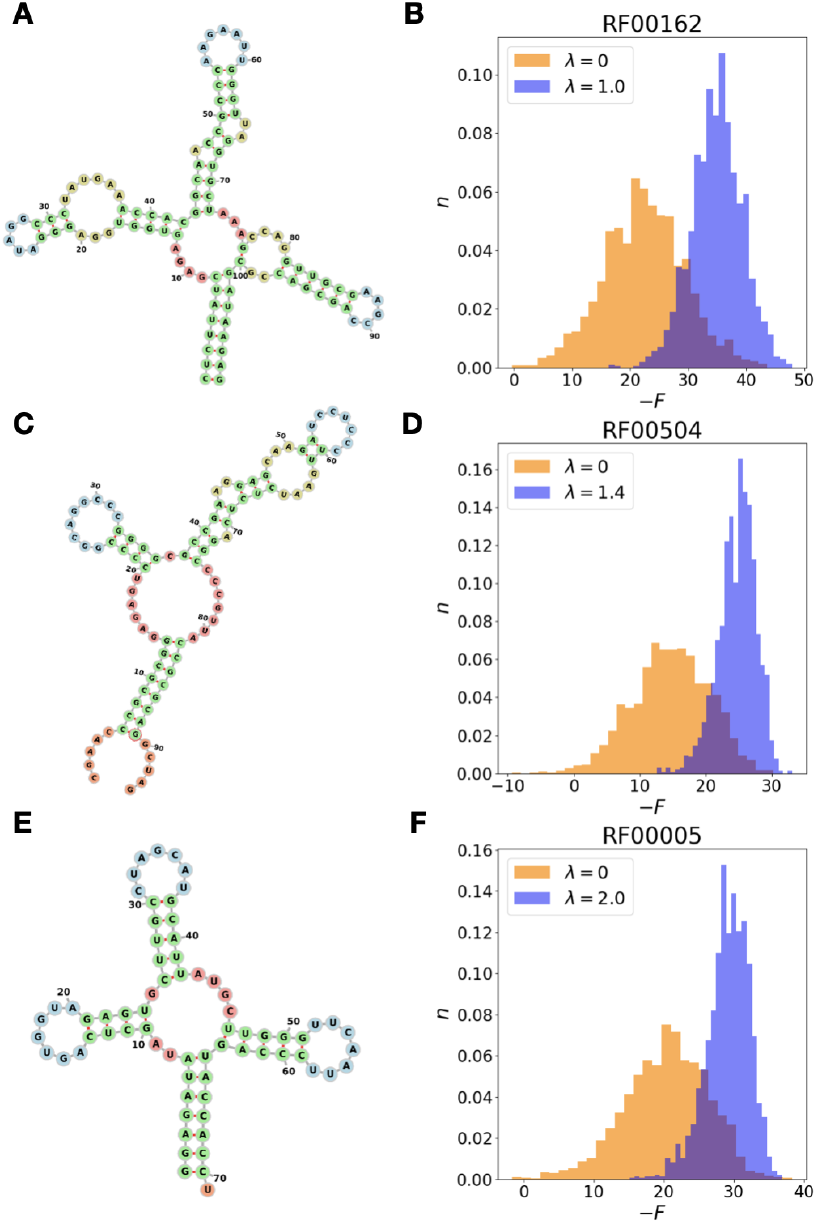
**A** Consensus secondary structure of the RF00162 RNA family (SAM riboswitch). **B** Distribution of the `RNAeval` proxy fitness for the RF00162 DCA model P^1^(<u>a</u>) (orange, λ = 0) and the reintegrated P^2^(<u>a</u>) model (blue, λ = 1). **C** Consensus secondary structure of the RF00504 RNA family (Glycine riboswitch). **D** Distribution of the `RNAeval` proxy fitness for the RF00504 DCA model P^1^(<u>a</u>) (orange, λ = 0, N = 2000) and the reintegrated P^2^(<u>a</u>) model (blue, λ = 2, N = 2000). **E** Consensus secondary structure of the RF00005 RNA family (tRNA). **F** Distribution of the `RNAeval` proxy fitness for the RF00005 DCA model P^1^(<u>a</u>) (orange, λ = 0, N = 2000) and the reintegrated P^2^(<u>a</u>) model (blue, λ = 1, N = 2000). The secondary structure diagrams were generated using the Forna software [17].

In addition to generating sequences with improved proxy fitness, another effect of the reintegration is that the DCA model score becomes a more reliable predictor of fitness, an in-depth analysis of this phenomenon is provided in the *Supplementary Section S2* and *Supplementary Figure S2*

### B. Reintegrating experimental activity of a protein family

Our case study for applying the reintegration procedure to proteins is the chorismate mutase (CM) enzyme, which plays an essential role in the biosynthesis of aromatic amino acids. This enzyme serves as an ideal setting to test our procedure because Russ et al. [1] have already trained a DCA model *P*^1^ on an MSA 𝒟_*N*_ of natural CM homologs, and they have experimentally tested the natural sequences of 𝒟_*N*_ as well as a dataset 𝒟_*T*_ of *P*^1^-designed CM variants using an *in vivo* growth assay (see *Materials and Methods*). As a result, we have access to experimentally labeled datasets indicating sequence functionality. Moreover, Russ et al. demonstrated that it is possible to train a simple Logistic Regression (LR) classifier on 𝒟_*N*_ to predict whether an artificial CM variant is functional or not. We can leverage these findings for our reintegration procedure.

Our approach begins by training an LR classifier using the labeled natural CM variants in 𝒟_*N*_ to predict experimental functionality. This classifier achieves an accuracy of approximately 80% in predicting the functionality of artificially generated sequences (cf. *Supplementary Section S3*), which is consistent with the results reported [1]. Since experimentally testing our artificial sequences is not feasible within this study, we will use this classifier to evaluate the performance of our reintegration procedure. Note that the labels used for training the classifier are not used in our reintegration procedure, but are complementary information exploitable for posterior sequence evaluation.

We first trained our initial DCA model *P*^1^(*<u>a</u>*) using the natural MSA 𝒟_*N*_ as training data, and the adabmDCA implementation of DCA [11]. From this model, we sampled the *P*^1^-dataset comprising 8000 artificial CM variants.

To train our reintegrated DCA model *P*^2^(*<u>a</u>*), we used dataset 𝒟_*T*_, which is experimentally labelled in [1], allowing us to avoid relying on proxy fitness measures. We chose again a binary adjustment function *w*(*<u>b</u>*) for all sequences in 𝒟_*T*_ :

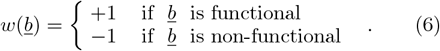

We set the reintegration strength parameter *λ* to 1, which is the highest value that converged within an acceptable time frame. The reintegration procedure is significantly slower for proteins compared to RNA, requiring 3 hours of runtime on an L4 GPU using the most advanced DCA GPU implementation available. Also here, we sampled from the resulting model a *P*^2^-dataset containing 8000 artificial CM sequences. Results for lower values of *λ* are provided in the *Supplementary Table S4*.

To assess the effectiveness of our reintegration procedure, we employed the LR classifier, and determined the percentage of variants predicted to be functional in both the *P*^1^- and the reintegrated *P*^2^-dataset.

The percentage of predicted functional variants increases from 39% to 68%, indicating a favorable outcome of the procedure. This comes at a moderate cost of reduced average sequence diversity, 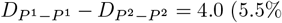 of 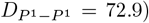, and the designs still retain a good level of diversity (SI). 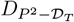 values and the histogram of distances between the *P* dataset and the closest reintegrated 𝒟_*T*_ are provided in the SI.

In the PCA plots shown in Fig. 4, we project the *P*^1^- and *P*^2^-generated datasets (colored) onto the first two principal components of the natural MSA 𝒟_*N*_ (grey). Notably, the *P*^2^ dataset avoids certain regions in PCA space that are occupied by sequences from both the natural dataset and the *P*^1^ model. To better understand this behavior, Fig. 5 projects the reintegration dataset 𝒟_*T*_ onto the same PCA, with functional sequences shown in green and non-functional ones in red. We observe that the regions avoided by the *P*^2^ dataset correspond to the non-functional areas in 𝒟_*T*_. These findings suggest that the observed reduction in diversity of sequences generated after reintegration results from the exclusion of non-functional regions, effectively refining the functional sequence space in the *P*^2^ model compared to the standard DCA model *P*^1^.

**FIG. 4:**
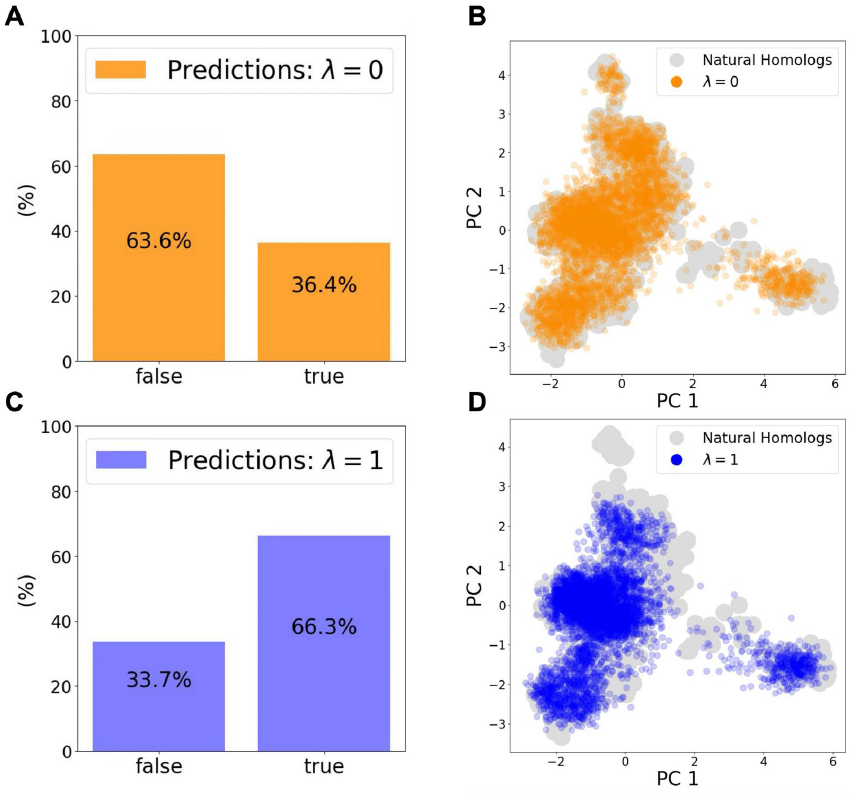
**A** Classifier predictions for protein functionality in the P^1^(<u>a</u>)-dataset (N = 8000): True indicates predicted functional, False indicates predicted non-functional. **B** PCA projection of the P^1^(<u>a</u>)-dataset (orange); natural PSA homologs are represented by the grey cloud. **C** Classifier predictions for protein functionality in the P^2^(<u>a</u>)-dataset (N = 8000): True indicates predicted functional, False indicates predicted non-functional. **D** PCA projection of the P^2^(<u>a</u>)-dataset (blue); natural PSA homologs are represented by the grey cloud.

**FIG. 5:**
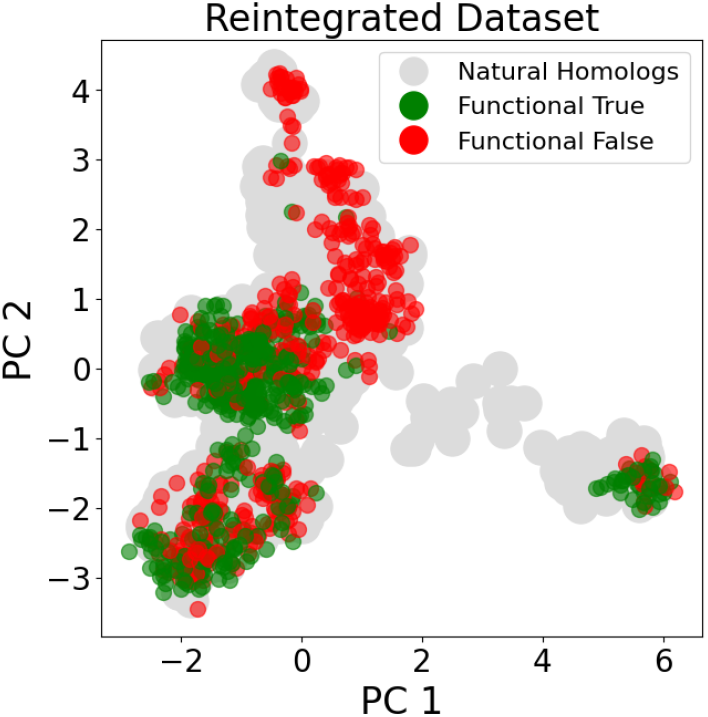
PCA projection of the experimentally labeled dataset D_T_ (N = 1003) of artificially generated chorismate mutase sequences. Functional sequences are shown in green, non-functional ones in red. The natural sequences in D_N_ (N = 1130) are shown in grey (note that not all of them are functional in the experimental test performed in E. coli).

### C. Experimental Validation on Group I Intron Ribozymes

Group I intron ribozymes are catalytic RNA molecules that possess the ability to self-splice, meaning they can excise themselves from precursor RNA transcripts without the assistance of proteins or additional enzymes. Our work leverages on that of Lambert et al. [2], who started with a reference wildtype, the *Azoarcus* group I intron ribozyme, and designed artificial mutations of this reference sequence using MSA-based generative models (DCA [15] and VAE [5]), structure based methods (Turner Model [13]) and combinations of the two. Through high-throughput assays, they tested these mutations for self-splicing-like activity and evaluated the percentage of active designs from each model at varying distances from the wild type. Here, we present experimental evidence supporting the presented reintegration method applied in this context.

We start with their DCA model, utilizing it as our non-reintegrated *P*^1^ model. This model was trained using as training set 𝒟_*N*_ an MSA of 817 group I introns aligned against the *Azoarcus* group I intron ribozyme (*L* = 197). For the training of our reintegrated models, we employ as 𝒟_*T*_ a subset of 14099 from their experimentally labeled dataset, which consists of 24071 mutations of the reference (see *Materials and Methods*).

An important detail is that the 𝒟_*T*_ in this case is significantly different from all the other reintegration instances in this study; all the tested designs are mutations of a single reference sequence, so their distribution in the sequence space is localized around one single point. Additionally, as expected, designs with fewer mutated residues generally exhibited higher activity than those with a greater number of mutations. Thus, a significant amount of information about sequence functionality contained in 𝒟_*T*_ is related to the trivial distance from the reference. Unlike all the other instances presented, where 𝒟_*T*_ designs are exclusively derived from the non-reintegrated model *P*^1^, in this case, they originate from all the different models tested in [2]. Details about 𝒟_*T*_ are provided in the *Supplementary Section S4* and *Supplementary Figure S3*. We implemented two reintegration strategies:

1. Standard Reintegration Procedure (REINT): All active 𝒟_*T*_ sequences 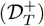 were reintegrated with a weight 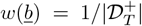, and all non-active 𝒟_*T*_ sequences 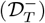 were reintegrated with 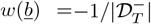.
2. Bin Sum Zero Reintegration (REINT BS0): 𝒟_*T*_ sequences were grouped into bins based on their mutational distance from the reference sequence, with each bin covering four mutational steps. For sequences in bin *i*, the weight *w*(*<u>b</u>*) was set to 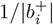 for active sequences, where 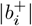 is the number of active sequences in the bin, and 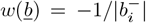 for non-active sequences, where 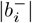 is the number of non-active sequences in the bin. This ensured that the sum of *w*(*<u>b</u>*) in each bin was zero, mitigating bias towards the *Azoarcus* reference sequence by balancing positive and negative signals at each distance.

For further details and the choice of the parameter *λ*, refer to the *Supplementary Section S4*.

The two models were trained using the same GPU DCA implementation used for the CM case.

From the two reintegrated models, we generated designs at various bins of mutational distance from the reference wildtype (Table 6). These designs were then experimentally tested for self-splicing activity using the same experimental assay that was used for the designs from the non-reintegrated *P*^1^ model [2]. Details about the experimental procedures and its comparability across different experimental istances are provided in the *Supplementary Section S4* and *Supplementary Figure S4, S5* and *Supplementary Table S5*.

The results of the experimental assays are displayed in Figure 6 and detailed in the *Supplementary Table S6*.

**FIG. 6:**
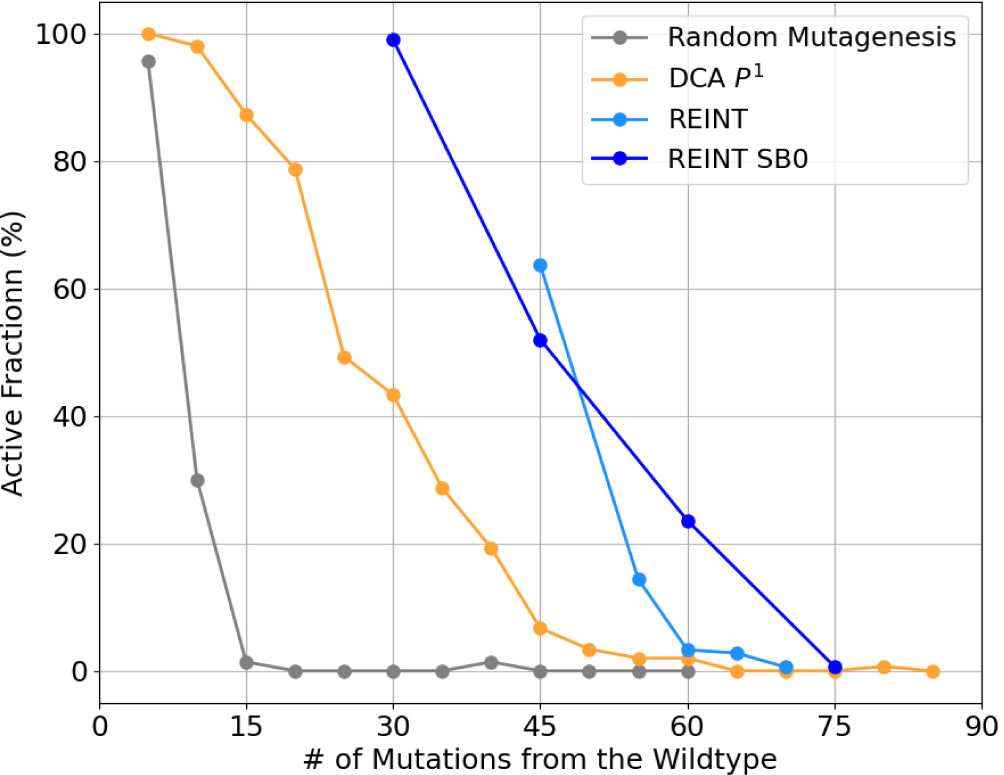
Active fraction of the generated designs as a function of the mutational distance from the reference sequence for the P^1^ DCA model and the two reintegrated models. The number of tested designs at each bin of distance can be found in Table II.

Both reintegrated models outperformed the non-reintegrated one. The REINT BS0 model maintained an active fraction of 23.6% at a mutational distance of 60 residues, compared to the non-reintegrated *P*^1^ model, which had a similar active fraction of 19.3% at 40 mutations but dropped to just 2.0% at 60 mutations. The REINT model successfully generated functional sequences at 65 mutations, further away than all the *P*^1^(*<u>a</u>*) tested in [2], where the furthest active seqwuence was found at 60 mutations.

This increase in performance came at the expense of generated sequence diversity, which was significantly more impacted than in the previous reintegration examples. This is most likely due to the highly localized nature of the reintegrated dataset 𝒟_*T*_. The intra-sample and 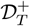 distance distributions for mutational distances of 45, 60 and 65 are shown in Table III, with the complete analysis provided in the *Supplementary Table S6* and *Supplementary Figure S6, S7*.

**TABLE II:** Number of tested designs of the two reintegrated models at various mutational distances. For the DCA P^1^ model, this number is 150 for each bin [2].

| Model | 30 | 45 | 55 | 60 | 65 | 70 | 75 |
| --- | --- | --- | --- | --- | --- | --- | --- |
| REINT | - | 80 | 180 | 180 | 180 | 180 | - |
| REINT BS0 | 100 | 150 | - | 275 | - | - | 275 |

**TABLE III:** Active Fraction, Average Intra-Dataset Distance 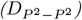, and Average Minimum Distance from the Positively Reintegrated Dataset 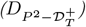 for mutational distances 60, 70, and 75 from the reference sequence.

| Model (Distance) | Active % | $D_{P^2-P^2}$ | $D_{P^2-\mathcal{D}_T^+}$ |
| --- | --- | --- | --- |
| DCA $P^1$ (45) | 6.7 | 51.3 | 31.8 |
| REINT (45) | 63.7 | 8.7 | 6.5 |
| REINT BS0 (45) | 52.0 | 10.0 | 7.6 |
| DCA $P^1$ (60) | 2.0 | 62.8 | 43.1 |
| REINT (60) | 3.3 | 12.5 | 18.6 |
| REINT BS0 (60) | 23.6 | 15.1 | 13.8 |
| DCA $P^1$ (65) | 0.0 | 70.4 | 50.3 |
| REINT (65) | 2.8 | 18.1 | 28.0 |

## III. DISCUSSION

Advances in high-throughput experimentation and machine learning are rapidly reshaping our ability to design functional biomolecular sequences. In this work, we have demonstrated that reintegrating experimental feedback into generative models markedly improves the reliability of predicted sequences. Our results show that even when the underlying mathematical model remains unchanged, the incorporation of well-characterized experimental data – including both functional and non-functional sequences – serves as a powerful corrective mechanism. Experimental tests therefore do not only validate predictions of models entirely trained on natural sequence data, but also guides the refinement of the modeled sequence space by penalizing non-functional regions and reinforcing areas associated with activity.

A key insight from our study is that enhanced performance can be achieved by training on more informative data rather than solely by increasing model complexity. The same DCA framework, when trained with a balanced integration of natural sequence data and experimental outcomes, produces a model that yields a significantly higher fraction of functional sequences. This observation underscores that the limitations of conventional generative models are not necessarily due to the inadequacy of their architectures, but rather stem from the sparsity and incomplete sampling inherent in natural sequence databases. By reintegrating experimental feedback, our method compensates for this deficiency, thereby sharpening the model’s discriminative power.

At the same time, our approach has inherent limitations. Because the reintegration procedure relies on experimental data that are necessarily sampled from regions already explored by nature, the model is not readily extended to completely novel areas of sequence space. In other words, while our method is highly effective at refining and navigating the known functional landscape, it remains dependent on the availability and quality of experimental measurements. This dependency emphasizes that the generation of truly novel sequences will continue to require a comprehensive experimental framework to guide and validate model predictions.

Looking ahead, the iterative interplay between experimental feedback and model refinement presents a promising route toward more accurate and targeted design strategies. Future studies could explore how successive rounds of data integration may gradually expand the functional sequence space, while also addressing potential trade-offs between accuracy and diversity. Ultimately, our results suggest that a synergistic integration of experimental and computational approaches can overcome the false-positive limitations of current generative models and pave the way for more reliable biomolecular design.

## IV. MATERIALS AND METHODS

In this section, we detail the methodological framework of our proposed reintegration method applied to the DCA generative model, as well as the data and methods employed for training and evaluating our models in a number of diverse RNA and protein families.

### A. Reintegration method for DCA models

The presented reintegration procedure is particularly well-suited for application to DCA models, as it does not require modifying their standard inference methods (differently from previous reintegration attempts [18]). In its conventional implementation, DCA assumes that the natural data distribution is described by a probability *P*^1^(*<u>a</u>*) in the form of a Potts model,

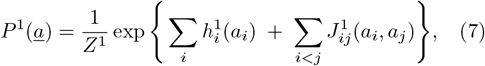

with *<u>a</u>* = (*a*_1_, …, *a*_*L*_) denoting, according to the problem under study, an aligned nucleotide or amino-acid sequence of length *L*. The optimal parameters {*h*^1^, *J* ^1^} for *P*^1^(*<u>a</u>*) are inferred via Maximum Likelihood Estimation (MLE) [12],

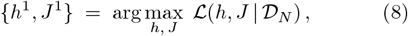

where 𝒟_*N*_ is an MSA of all sequenced homologs of the family under consideration.

We omit here the standard sequence reweighting procedure [19] commonly used in DCA, to simplify notation, but it can be included straightforwardly. Our publicly available implementation does contain it. The reintegration data consist of a second alignment, 𝒟_*T*_, of experimentally tested sequences, together with experimental outcomes encoded by the adjustment weight *w*(*<u>b</u>*). The reintegrated model *P*^2^(*<u>a</u>*) remains in the Potts-model form,

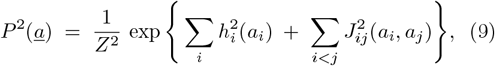

but with updated parameters {*h*^2^, *J* ^2^} chosen to maximize the new objective function

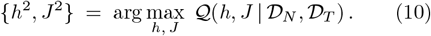

In standard DCA inference, the MLE conditions require the one-point marginals 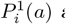 and two-point marginals 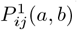 of *P*^1^(*<u>a</u>*) to match the empirical frequencies *f*_*i*_(*a*) and *f*_*ij*_(*a, b*) observed in 𝒟_*N*_ :

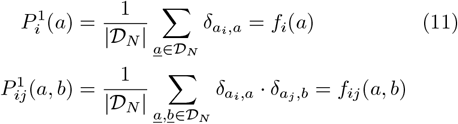

where *δ*_*a,b*_ is the Kronecker delta.

By contrast, maximizing 𝒬 imposes choosing {*h*^2^, *J* ^2^} so that the one-point and two-point marginals of *P*^2^(*<u>a</u>*) match the 𝒟_*T*_ -corrected frequencies 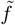 (see *Supplementary Section S1*):

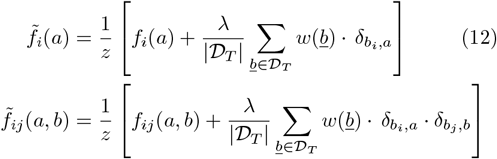

where

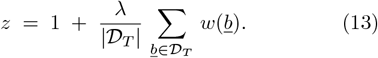

Hence, the maximization condition becomes

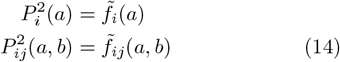

for all *i, j* and all *a, b*. Thus, any standard DCA inference method may be used simply by replacing the original empirical frequencies *f* with the corrected frequencies 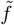 from Eq. (12). Consequently, implementing reintegration in a DCA pipeline requires only altering the frequency targets used during parameter inference.

It is important to note, however, that negative values of *w*(*<u>b</u>*) (i.e. experimentally tested non-functional sequences) can break the convexity of the problem, removing guaranteed uniqueness or even existence of a solution. Additionally, the corrected frequencies 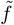 may sometimes lie outside the interval [0, 1]. As a result, convergence to a consistent DCA model is not strictly guaranteed, especially for strong reintegration strengths *λ*. Nonetheless, in all of our applications, we found that moderate settings of *λ* do converge reliably.

### B. Data, RNA fitness proxy & Group I intron experimental activity

#### 1. RFAM sequence data

− The RNA datasets used in this paper are MSA of three RNA families. These families are the tRNA family RF00005 (number of sequences |𝒟_*N*_| = 30000, number of residues *L* = 71), the SAM riboswitch family RF0162 (|𝒟_*N*_| = 6113, *L* = 108), and the Glycine riboswitch family RF0504 (|𝒟_*N*_| = 4600, *L* = 94). For all three RNA families, the corresponding consensus secondary structure is available from the Rfam database [7]. The Datasets are taken from [15].

#### 2. RNA fitness proxy

− To study the reintegration procedure, we need a fast and reliable way to label our datasets. Performing experiments on RNA sequence functionality is both expensive and time-consuming, so we initially used computational tools as fitness proxies to label our datasets. Since we have access to the RNA families’ consensus secondary structures, we employed the `RNAeval` function provided by the Vienna Package [14]. `RNAeval` calculates the thermodynamic free energy *F* (using the Turner 2004 energy model [13]) of an RNA sequence folded onto a given secondary structure.

Our underlying assumption is that well-generated sequences will, on average, exhibit low free energy *F* when folded onto the family’s consensus structure, indicating structural stability. In contrast, poorly generated sequences may lack structural stability or fold into alternative structures, and are expected to have a higher free energy. Therefore, we will use −*F* as a fitness proxy. This approach leverages the established structure/function relationship in RNA [20, 21], and enables us to computationally label sequences for a given target secondary structure.

#### 3. Group I Intron Ribozymes Data

− The Group I intron datasets utilized in this paper are taken from the study by Lambert et al. [2]. These datasets consist of multiple sequence alignments (MSAs) of Group I intron ribozymes. The natural data *D*_*N*_ consists of an MSA of *N* = 817 natural Group I introns aligned against the reference sequence, the Azoarcus Group I intron (numbeer of residues *L* = 197). The reintegration dataset *D*_*T*_ consists of a subset of 14099 out of the original 24071 experimentally labeled sequences. Specifically, *D*_*T*_ includes the sequences with mutational distance from 3 to 60, excluding those near the activity threshold. Sequences with mutational distances exceeding 60 were not reintegrated due to the unreliable detection of activity signals beyond this range (see *Supplementary Section S4*).

#### 4. Group I Experimental Activity

− To experimentally validate the reintegration methods, we conducted experiments on artificially designed Group I intron ribozymes. The experimental activity was determined using the same methodology as the tested ribozymes in Lambert et al. [2]. The experimental assay consists of a high-throughput screening of self-splicing catalytic activity, details are provided in the *Supplementary Section S4*.

#### 5. Protein data

The protein datasets used in this paper are taken from the study by Russ et al. [1]. These datasets consist of MSAs of the chorismate mutase (CM) enzyme. 𝒟_*N*_ consists of the MSA of the natural CM homologs (number of sequences *N* = 1130, number of residues *L* = 96) and the reintegration dataset 𝒟_*T*_ is the alignment (*N* = 1003, *L* = 96) of DCA-generated artificial CM variants. In [1], both datasets are labeled based on experimental testing: all protein sequences were expressed in genetically engineered *E. coli* strains, each modified to produce one of the CMs from 𝒟_*N*_ or 𝒟_*T*_ instead of their natural wildtype variants. These *E. coli* strains were then tested for growth under selective conditions; sequences enabling *E. coli* growth were labeled as functional, whereas those that did not were labeled as non-functional. It is noteworthy that many natural sequences in 𝒟_*N*_ were non-functional in *E. coli* under the experiment’s conditions: the natural CM homologs have undergone evolutionary adaptations specific to their native hosts, and may fail to provide growth in the *E. coli* environment.

## DATA AND CODE AVAILABILITY

All data and code used in this study are publicly available at Zenodo: 10.5281/zenodo.15115193.

## ACKNOWLEDGEMENTS

We acknowledge helpful discussions with Camille Lambert, Roberto Netti, Vaitea Opuu, Andrea Pagnani, Lorenzo Rosset and Francesco Zamponi during the preparation of this work, and we thank particularly Lorenzo Rosset for providing his GPU-based adabmDCA implementation prior to publication, and Vaitea Opuu for his help with the initial analysis of the experimental data using the scripts of [2].

