## Supplemental information for "Integrating experimental feedback improves generative models for biological sequences"

Francesco Calvanese,[1,2,*] Giovanni Peinetti,[1,*] Polina Pavlinova,[2] Philippe Nghe,[2] and Martin Weigt[1]
[1]*Sorbonne Université, CNRS, Dept. of Computational, Quantitative and Synthetic Biology - CQSB, Paris, France*
[2]*Laboratoire de Biophysique et Evolution, CNRS-ESPCI, PSL University, Paris, France*

### S1. DCA ANALYTICAL COMPUTATIONS

$P^2(\underline{a})$ is a Boltzmann distribution over the sequence space,

$$P^2(\underline{a}) = \frac{1}{Z^2} \exp\left\{ -H^2(\vec{a})\right\} \ , \tag{S1}$$

defined via the Potts Hamiltonian $H^2(\vec{a})$

$$H^2(\underline{a}) = \sum_i h_i^2(a_i) + \sum_{i<j} J_{ij}^2(a_i, a_j) \ . \tag{S2}$$

The optimal parameters $\{h^2, J^2\}$ are obtained via

$$\{h^2, J^2\} = \underset{h,J}{\operatorname{argmax}}\ \mathcal{Q}(h, J|\mathcal{D}_N, \mathcal{D}_T) \tag{S3}$$

as arguments of the maximum of the new objective function

$$Q(h, J|\mathcal{D}_N, \mathcal{D}_T) = \frac{1}{|\mathcal{D}_N|} \sum_{\underline{a}\in\mathcal{D}_N} \ln P^2(\underline{a}|h, J) + \frac{\lambda}{|\mathcal{D}_T|} \sum_{\underline{b}\in\mathcal{D}_T} w(\underline{b}) \cdot \ln P^2(\underline{b}|h, J) \ . \tag{S4}$$

It is possible to exploit Eq. (S1) to write :

$$\mathcal{Q} = -\frac{1}{|\mathcal{D}_N|} \sum_{\underline{a}\in\mathcal{D}_N} H^2(\underline{a}) - \log Z^2 - \frac{\lambda}{|\mathcal{D}_T|} \sum_{\underline{b}\in\mathcal{D}_T} H^2(\underline{b}) \cdot w(\vec{b}) - \frac{\lambda}{|\mathcal{D}_T|} \log Z^2 \sum_{\underline{b}\in\mathcal{D}_T} w(\underline{b}) \ .$$

The aim is to analytically maximize $Q(h, J|\mathcal{D}_N, \mathcal{D}_T)$ to find $\{h^2, J^2\}$. To this end, we need to compute partial derivatives of $Q$ with respect to fields $h_i^2(a)$ and couplings $J_{ij}^2(a, b)$. Proceeding term by term we find

$$\begin{aligned} -\frac{1}{|\mathcal{D}_N|} \sum_{\underline{a}\in\mathcal{D}_N} \frac{\partial H^2(\underline{a})}{\partial h_i^2(a)} &= -\frac{1}{|\mathcal{D}_N|} \sum_{\underline{a}\in\mathcal{D}_N} \frac{\partial\left(\sum_j h_j^2(a_j)\right)}{\partial h_i^2(a)} \\ &= -\frac{1}{|\mathcal{D}_N|} \sum_{\underline{a}\in\mathcal{D}_N} \delta_{a_i,a} \\ &= f_i(a) \ . \end{aligned} \tag{S5}$$

Similarly we obtain

$$-\frac{1}{|\mathcal{D}_N|} \sum_{\underline{a}\in\mathcal{D}_N} \frac{\partial H^2(\underline{a})}{\partial J_{ij}^2(a, b)} = f_{ij}(a, b) \ .$$

---

* These authors contributed equally to this work.

The derivatives of the logarithm of partition function can be computed exploiting the same reasoning of Eq. (S5),

$$
\begin{aligned}
-\frac{\partial \log Z^2}{\partial h_i^2(a)} &= \frac{1}{Z^2}\frac{\partial Z^2}{\partial h_i^2(a)} = \frac{1}{Z^2}\sum_{\underline{a}} \frac{\partial}{\partial h_i^2(a)} \exp\big\{-H^2(\underline{a})\big\} \qquad \text{(S6)}\\
&= \frac{1}{Z^2}\sum_{\underline{a}} \delta_{a_i,a}\cdot \exp\big\{-H^2(\underline{a})\big\}\\
&= \sum_{\underline{a}} \delta_{a_i,a}\cdot P^2(\vec{a}) = P_i^2(a)\ .
\end{aligned}
$$

Similarly we obtain

$$
-\frac{\partial \log Z^2}{\partial J_{ij}^2(a,b)} = P_{ij}^2(a,b)\ .
$$

Finally, using again Eq. (S5), the derivatives of terms involving the adjustment function $w(\vec{b})$ can be computed,

$$
\begin{aligned}
-\sum_{\underline{b}\in\mathcal{D}_T} \frac{\partial H^2(\underline{b})\cdot w(\underline{b})}{\partial h_i^2(a)} &= \sum_{\underline{b}\in\mathcal{D}_T} \delta_{b_i,a}\cdot w(\underline{b}) \qquad \text{(S7)}\\
-\sum_{\underline{b}\in\mathcal{D}_T} \frac{\partial H^2(\underline{b})\cdot w(\underline{b})}{\partial J_{ij}^2(a,b)} &= \sum_{\underline{b}\in\mathcal{D}_T} \delta_{b_i,a}\cdot\delta_{b_j,b}\cdot w(\underline{b})
\end{aligned}
$$

Rearranging terms, the following equations for the first and second moment of $P^2(\vec{a})$ are found:

$$
\begin{aligned}
P_i^2(a) &= \frac{1}{z}\left[f_i(a) + \frac{\lambda}{|\mathcal{D}_T|}\sum_{\underline{b}\in\mathcal{D}_T} w(\underline{b})\cdot\delta_{b_i,a}\right] = \tilde{f}_i(a) \qquad \text{(S8)}\\
P_{ij}^2(a,b) &= \frac{1}{z}\left[f_{ij}(a,b) + \frac{\lambda}{|\mathcal{D}_T|}\sum_{\underline{b}\in\mathcal{D}_T} w(\underline{b})\cdot\delta_{b_i,a}\cdot\delta_{b_j,b}\right] = \tilde{f}_{ij}(a,b)
\end{aligned}
$$

with normalization

$$
z = 1 + \frac{\lambda}{|\mathcal{D}_T|}\sum_{\vec{b}\in\mathcal{D}_T} w(\underline{b})\ . \qquad \text{(S9)}
$$

The model training can be performed using all the standard DCA training techniques, using the adjusted frequencies $\tilde{f}_i$ and $\tilde{f}_{ij}$ in Eq. S8 as targets for the model's marginals instead of the empirical MSA frequencies $f_i$ and $f_{ij}$.

### S2. RNA RFAM FAMILIES

#### S2.1. Edge Activation DCA Training

For each RFAM family, we used the Edge Activation DCA (eaDCA) algorithm from [1] to train a DCA model $P^1(\underline{a})$ on the MSA of the RNA family. We used 8000 chains for training and a pseudocount of 0.05. The empirical frequencies $f_i(a)$ and $f_{ij}(a,b)$ are computed along with the correlation matrix $C_{ij}^{emp}(a,b) = f_{ij}(a,b) - f_i(a)f_j(b)$. Training stops when the Pearson correlation between $C_{ij}^{emp}(a,b)$ and $C_{ij}^{train}(a,b)$ reaches 0.95.
Once the model is learned, we sample artificial sequences with Gibbs Sampling, forcing the obtained sequences to have no gaps. This is done to ensure that the `RNAeval` proxy fitness, $-F$ is comparable across different sequences. A dataset of 2000 artificial sequences was sampled from each model. These sequences, along with a `RNAeval` proxy fitness $-F$ for each of them, serve as Reintegration Dataset $\mathcal{D}_T$ for our reintegration procedure.
Subsequently, we computed the effective frequencies using Equation S8 to train the reintegrated model $P^2(\underline{a})$. Depending on the value of $\lambda$, the convergence of the reintegrated model's training is not always guaranteed. To facilitate convergence of the eaDCA procedure, we set any negative effective frequencies to zero and enforced a constraint to prevent the same edge from being activated more than five times. All other training settings were kept identical to those used for training the non-reintegrated $P^1(\underline{a})$ model.

#### S2.2. Results for different values of $\lambda$

We tested the reintegration procedure at different values of $\lambda$, starting with $\lambda = 0$ and increasing $\lambda$ by 0.1 at each step. The maximum $\lambda$ tested corresponds to the largest value below 2 for which our new model converges within $10^4$ steps using the eaDCA algorithm. For RF00504, the resulting interval of tested $\lambda$ values is $[0.1, 1.4]$, for RF00162 $[0.1, 1]$, and for RF00005 $[0.1 : 2]$. The complete results of these analyses are presented for all three families in Tables S1, S2, and S3.
Furthermore, we provide, for each family, the histograms of `RNAeval` [2] proxy fitness $-F$ for $\lambda = 0.1$, 0.5 and $\lambda = \lambda_{max}$. These can be found in Figure 1.

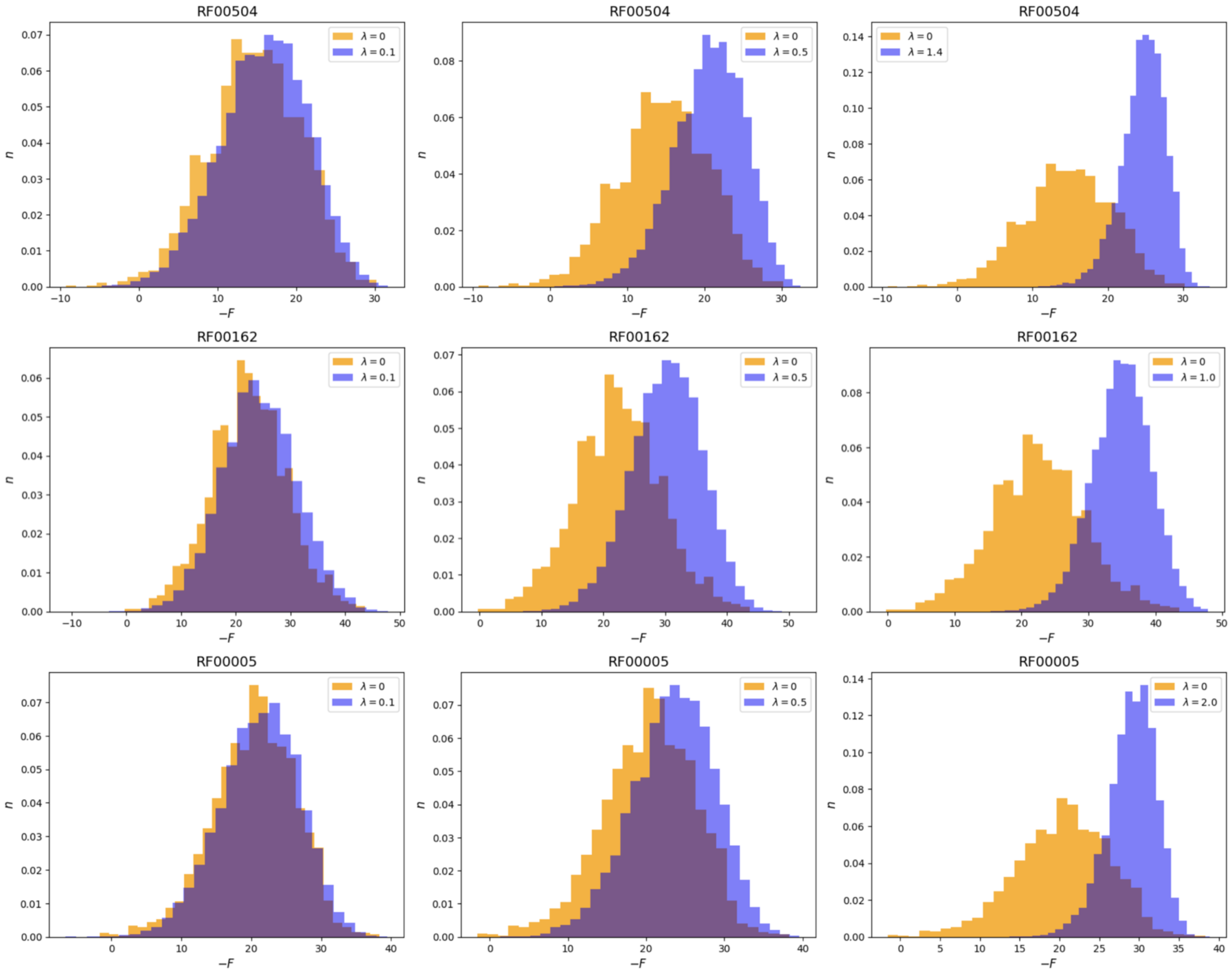


FIG. S1:
*Distribution of the* `RNAeval` *proxy fitness for the RF00504 DCA model $P^1(\underline{a})$ (orange, $\lambda = 0$) and the reintegrated $P^2(\underline{a})$ model (blue, $\lambda = 0.1$, $\lambda = 0.5$, $\lambda =$ 1.4)*
*Distribution of the* `RNAeval` *proxy fitness for the RF00162 DCA model $P^1(\underline{a})$ (orange, $\lambda = 0$) and the reintegrated $P^2(\underline{a})$ model (blue, $\lambda = 0.1$, $\lambda = 0.5$, $\lambda =$ 1.0)*
*Distribution of the* `RNAeval` *proxy fitness for the RF00005 DCA model $P^1(\underline{a})$ (orange, $\lambda = 0$) and the reintegrated $P^2(\underline{a})$ model (blue, $\lambda = 0.1$, $\lambda = 0.1$, $\lambda =$ 2.0)*

| $\lambda$ value | TP | $-\tilde{F}$ | $S$ | $D_{P^2-P^2}$ | $D_{P^2-\mathcal{D}_T^+}$ | $D_{P^2-\mathcal{D}_T^-}$ |
|---|---|---|---|---|---|---|
| 0.0 | 51.6% | 14.7 | 61.9 | 43.3 | – | – |
| 0.1 | 60.3% | $15.9 \pm 0.1$ | 63.8 | $42.8 \pm 0.1$ | $26.0 \pm 0.1$ | $27.8 \pm 0.1$ |
| 0.2 | 69.6% | $17.2 \pm 0.1$ | 63.3 | $41.2 \pm 0.1$ | $25.0 \pm 0.1$ | $27.5 \pm 0.1$ |
| 0.3 | 80.6% | $18.9 \pm 0.1$ | 62.9 | $39.4 \pm 0.1$ | $24.2 \pm 0.1$ | $27.3 \pm 0.1$ |
| 0.4 | 84.5% | $19.6 \pm 0.1$ | 62.6 | $38.5 \pm 0.1$ | $23.8 \pm 0.1$ | $27.2 \pm 0.1$ |
| 0.5 | 89.6% | $20.6 \pm 0.1$ | 61.2 | $37.3 \pm 0.1$ | $23.3 \pm 0.1$ | $27.1 \pm 0.1$ |
| 0.6 | 93.3% | $21.7 \pm 0.1$ | 60.6 | $35.6 \pm 0.1$ | $22.7 \pm 0.1$ | $26.8 \pm 0.1$ |
| 0.7 | 96.1% | $22.5 \pm 0.1$ | 60.1 | $34.7 \pm 0.2$ | $22.2 \pm 0.1$ | $26.6 \pm 0.1$ |
| 0.8 | 95.8% | $22.4 \pm 0.1$ | 58.7 | $34.1 \pm 0.1$ | $21.9 \pm 0.1$ | $26.4 \pm 0.1$ |
| 0.9 | 97.6% | $23.1 \pm 0.1$ | 57.7 | $33.7 \pm 0.1$ | $21.7 \pm 0.1$ | $26.4 \pm 0.1$ |
| 1.0 | 98.6% | $23.7 \pm 0.1$ | 55.9 | $33.2 \pm 0.1$ | $21.4 \pm 0.1$ | $26.2 \pm 0.1$ |
| 1.1 | 98.7% | $23.8 \pm 0.1$ | 53.2 | $32.7 \pm 0.1$ | $21.2 \pm 0.1$ | $26.1 \pm 0.1$ |
| 1.2 | 99.3% | $24.3 \pm 0.1$ | 51.6 | $32.5 \pm 0.1$ | $21.0 \pm 0.1$ | $26.1 \pm 0.1$ |
| 1.3 | 99.4% | $24.5 \pm 0.1$ | 48.3 | $32.0 \pm 0.1$ | $20.6 \pm 0.1$ | $25.9 \pm 0.1$ |
| 1.4 | 99.7% | $24.7 \pm 0.1$ | 43.3 | $31.6 \pm 0.1$ | $20.3 \pm 0.1$ | $25.7 \pm 0.1$ |

TABLE S1: RF504

| $\lambda$ value | TP | $-\tilde{F}$ | $S$ | $D_{P^2-P^2}$ | $D_{P^2-\mathcal{D}_T^+}$ | $D_{P^2-\mathcal{D}_T^-}$ |
|---|---|---|---|---|---|---|
| 0.0 | 49.9% | 22.4 | 63.5 | 46.0 | – | – |
| 0.1 | 60.4% | $24.3 \pm 0.2$ | 65.3 | $45.8 \pm 0.1$ | $28.1 \pm 0.1$ | $29.6 \pm 0.1$ |
| 0.2 | 69.5% | $25.9 \pm 0.1$ | 65.6 | $44.9 \pm 0.1$ | $27.7 \pm 0.1$ | $29.8 \pm 0.1$ |
| 0.3 | 80.0% | $27.7 \pm 0.1$ | 64.7 | $43.8 \pm 0.2$ | $27.2 \pm 0.1$ | $30.0 \pm 0.1$ |
| 0.4 | 85.2% | $28.9 \pm 0.1$ | 63.9 | $42.9 \pm 0.1$ | $27.0 \pm 0.1$ | $30.3 \pm 0.1$ |
| 0.5 | 91.3% | $30.2 \pm 0.2$ | 63.2 | $42.2 \pm 0.1$ | $26.7 \pm 0.1$ | $30.5 \pm 0.1$ |
| 0.6 | 94.7% | $31.3 \pm 0.1$ | 62.3 | $41.6 \pm 0.1$ | $26.4 \pm 0.1$ | $30.6 \pm 0.1$ |
| 0.7 | 96.6% | $32.2 \pm 0.1$ | 61.4 | $40.9 \pm 0.1$ | $26.2 \pm 0.1$ | $30.9 \pm 0.1$ |
| 0.8 | 98.0% | $33.0 \pm 0.1$ | 59.4 | $40.3 \pm 0.1$ | $25.9 \pm 0.1$ | $31.0 \pm 0.1$ |
| 0.9 | 98.9% | $33.8 \pm 0.1$ | 57.1 | $39.5 \pm 0.1$ | $25.6 \pm 0.1$ | $31.0 \pm 0.1$ |
| 1.0 | 99.4% | $34.9 \pm 0.1$ | 53.3 | $39.7 \pm 0.1$ | $25.3 \pm 0.1$ | $31.8 \pm 0.1$ |

TABLE S2: RF162

| $\lambda$ value | TP | $-\tilde{F}$ | $S$ | $D_{P^2-P^2}$ | $D_{P^2-\mathcal{D}_T^+}$ | $D_{P^2-\mathcal{D}_T^-}$ |
|---|---|---|---|---|---|---|
| 0.0 | 52.3% | 20.4 | 50.3 | 36.3 | – | – |
| 0.1 | 57.7% | $21.2 \pm 0.2$ | 51.2 | $36.1 \pm 0.1$ | $20.7 \pm 0.1$ | $22.2 \pm 0.1$ |
| 0.2 | 61.6% | $21.7 \pm 0.1$ | 50.9 | $35.8 \pm 0.1$ | $20.5 \pm 0.1$ | $22.1 \pm 0.1$ |
| 0.3 | 65.6% | $22.3 \pm 0.1$ | 50.2 | $35.7 \pm 0.1$ | $20.4 \pm 0.1$ | $22.1 \pm 0.1$ |
| 0.4 | 69.2% | $22.9 \pm 0.2$ | 50.0 | $35.5 \pm 0.1$ | $20.2 \pm 0.1$ | $22.0 \pm 0.1$ |
| 0.5 | 73.3% | $23.5 \pm 0.1$ | 49.5 | $35.3 \pm 0.1$ | $20.0 \pm 0.1$ | $22.0 \pm 0.1$ |
| 0.6 | 76.4% | $23.9 \pm 0.2$ | 49.0 | $35.0 \pm 0.1$ | $19.8 \pm 0.1$ | $21.9 \pm 0.1$ |
| 0.7 | 80.1% | $24.4 \pm 0.2$ | 48.9 | $34.8 \pm 0.1$ | $19.7 \pm 0.1$ | $21.9 \pm 0.1$ |
| 0.8 | 83.3% | $24.9 \pm 0.1$ | 47.6 | $34.6 \pm 0.1$ | $19.5 \pm 0.1$ | $21.9 \pm 0.1$ |
| 0.9 | 86.0% | $25.4 \pm 0.1$ | 47.6 | $34.3 \pm 0.1$ | $19.3 \pm 0.1$ | $21.9 \pm 0.1$ |
| 1.0 | 88.6% | $25.7 \pm 0.1$ | 46.6 | $34.2 \pm 0.1$ | $19.3 \pm 0.1$ | $21.9 \pm 0.1$ |
| 1.1 | 91.2% | $26.2 \pm 0.1$ | 46.1 | $34.1 \pm 0.1$ | $19.1 \pm 0.1$ | $21.9 \pm 0.1$ |
| 1.2 | 93.9% | $26.8 \pm 0.1$ | 45.3 | $33.6 \pm 0.1$ | $18.9 \pm 0.1$ | $21.7 \pm 0.1$ |
| 1.3 | 94.7% | $27.0 \pm 0.1$ | 44.4 | $33.5 \pm 0.1$ | $18.8 \pm 0.1$ | $21.7 \pm 0.1$ |
| 1.4 | 96.0% | $27.4 \pm 0.1$ | 43.8 | $33.4 \pm 0.1$ | $18.7 \pm 0.1$ | $21.7 \pm 0.1$ |
| 1.5 | 97.6% | $27.8 \pm 0.1$ | 42.8 | $33.3 \pm 0.1$ | $18.6 \pm 0.1$ | $21.8 \pm 0.1$ |
| 1.6 | 98.1% | $28.2 \pm 0.1$ | 41.5 | $32.9 \pm 0.1$ | $18.4 \pm 0.1$ | $21.7 \pm 0.1$ |
| 1.7 | 98.4% | $28.3 \pm 0.1$ | 39.8 | $32.8 \pm 0.1$ | $18.3 \pm 0.1$ | $21.7 \pm 0.1$ |
| 1.8 | 98.8% | $28.5 \pm 0.1$ | 38.8 | $32.7 \pm 0.1$ | $18.2 \pm 0.1$ | $21.8 \pm 0.1$ |
| 1.9 | 99.0% | $28.8 \pm 0.1$ | 36.5 | $32.3 \pm 0.1$ | $18.0 \pm 0.1$ | $21.7 \pm 0.1$ |
| 2.0 | 99.1% | $29.1 \pm 0.1$ | 33.7 | $32.3 \pm 0.1$ | $18.0 \pm 0.1$ | $21.8 \pm 0.1$ |

TABLE S3: RF005

#### S2.3. Fitness landscape prediction

Previous studies [3, 4] suggest there should be an anti-correlation between the energy assigned by the model and the actual fitness of a sequence. In this section, we examine whether our experiment-informed reintegrated model improves its ability to predict sequence fitness.
Using each of the RNA families RF00504, RF00005, and RF00162, we generated datasets $D_{\text{global}}$, consisting of 2000 independent sequences sampled from the non-reintegrated $P^1(\underline{a})$ model with Gibbs Sampling.
Since $P^1(\underline{a})$ was trained on the Natural MSA, these sequences display diversity similar to that of their respective natural RNA families and are widely distributed across the sequence space (Table S1, S2, S3).
For each sequence in $D_{\text{global}}$, we calculated the energies of both the $P^1(\underline{a})$ and $P^2(\underline{a})$ models and measured their correlation with the proxy fitness $-F$. Across all three RNA families, we observe a substantial increase in the correlation between model energy and proxy fitness after reintegration. This improvement depends on the reintegration strength parameter $\lambda$, with higher $\lambda$ values leading to stronger correlation enhancements. This can be seen in Figure S5.

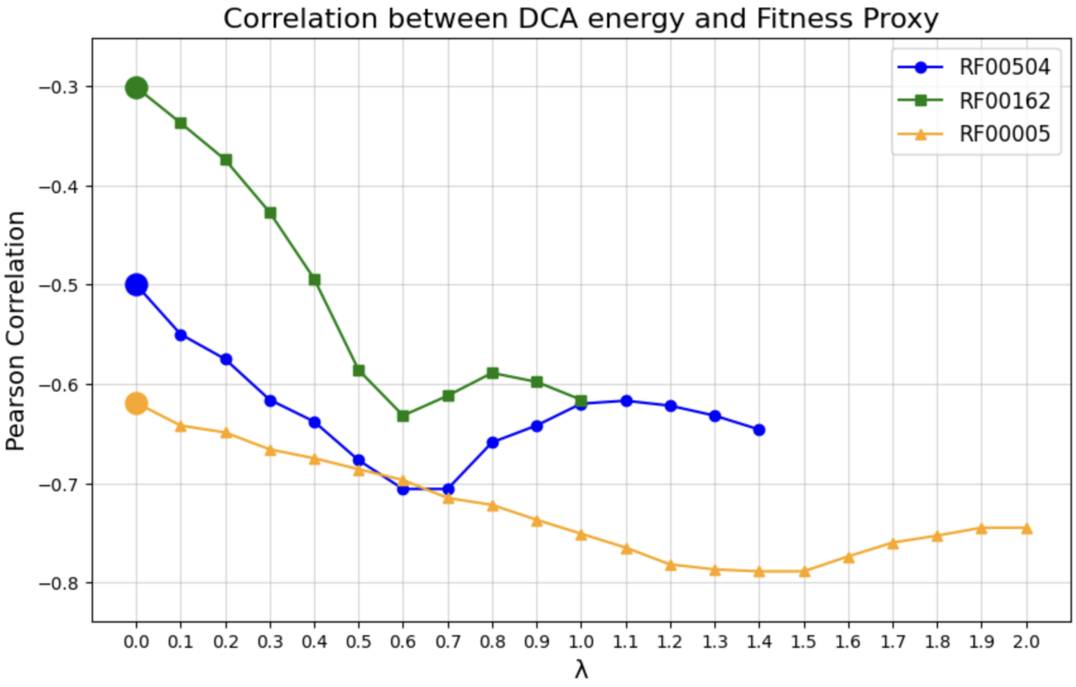


FIG. S2: *Correlation between DCA energy and fitness proxy for three RNA families (RF00504, RF00162, RF00005) across the considered range of $\lambda$ values. At $\lambda = 0$ (without reintegration), the starting points for each family are highlighted.*

### S3. CHORISMATE MUTASE

#### S3.1. Classifier prediction for CM

As described in [5], a logistic regression classifier is used to predict sequence functionality. The training set for the logistic regression consists of the experimentally labeled natural homologs ($\mathcal{D}_N$) published in [5]. Each sequence is assigned a binary label $x$, where $x = 1$ denotes a sequence found to be functional under experimental assay conditions, and $x = 0$ corresponds to an experimentally non-functional sequence. The predicted probability of a sequence $\underline{a}$ to be functional is given by

$$P(x = 1 \,|\, \underline{a}) \sim \exp\left\{ g + \sum_{i=1}^{L} K_i(a_i) \right\},$$

where $g$ represents a bias term, and $K_i(a_i)$ links the functionality $x$ to the specific amino acid $a_i$ at position $i$.
The accuracy of this classifier, when tested on the $\mathcal{D}_T$ protein dataset, is about 80%.

#### S3.2. Training Procedure and Results for different values of $\lambda$

To apply the reintegration procedure on the Chorismate Mutase protein family, we used the Adaptive Boltzmann Machine DCA (adabmDCA) algorithm to train a DCA model $P^2(\underline{a})$ using the effective frequencies computed at different values of $\lambda$. Depending on the value of $\lambda$, the convergence of the reintegrated model's training is not always guaranteed. To facilitate convergence of the adabmDCA procedure, we set any negative effective frequencies to zero. All other training settings were kept identical to those used for training the non-reintegrated $P^1(\underline{a})$ model. The training was performed with 10,000 Monte Carlo chains, 10 sweeps per gradient update, and a learning rate of 0.05. Training stopped once the Pearson correlation between the empirical and training correlation matrices, $C_{ij}^{emp}(a,b)$ and $C_{ij}^{train}(a,b)$, reached 0.95. The results of the reintegration procedure for $\lambda = \{0.25, 0.5, 0.75, 1.0\}$ are shown in Table S4.

| $\lambda$ value | Working (%) | $D_{P^2-P^2}$ | $D_{P^2-\mathcal{D}_T^+}$ | $D_{P^2-\mathcal{D}_T^-}$ |
|---|---|---|---|---|
| $\lambda = 0$ | 36.4 | 72.9 | – | – |
| $\lambda = 0.25$ | 43.6 | 72.0 | 49.4 | 50.9 |
| $\lambda = 0.5$ | 51.5 | 70.9 | 46.9 | 50.0 |
| $\lambda = 0.75$ | 59.5 | 69.9 | 43.9 | 48.5 |
| $\lambda = 1$ | 66.3 | 68.9 | 40.4 | 46.6 |

TABLE S4: *Percentage of sequences classified as functional, average intra-dataset distance ($D_{P^2-P^2}$), and average minimum distance from the positively reintegrated dataset ($D_{P^2-\mathcal{D}_T^+}$) for samples coming from reintegrated models trained for Chorismate Mutases at different values of the reintegration strength $\lambda$.*

### S4. GROUP I INTRON RYBOZIMES

#### S4.1. Reintegration $\mathcal{D}_T$ Dataset

The ribozyme reintegration dataset $\mathcal{D}_T$ consists of 14099 experimentally annotated sequences out of the 24071 [6] tested ones, all of length 197. Fig. S3 illustrates the experimentally determined activity and mutational distance from the Azoarcus reference sequence for all tested sequences in [6]. Positively reintegrated sequences are shown in green, while negatively reintegrated sequences are shown in red. Reintegrated sequences have between 4 and 60 mutations from wildtype, beyond which no reliable activity signal was detected. We also excluded sequences in the direct vicinity of the activity threshold from the reintegration set, to avoid noisy activity annotations to be reintegrated.

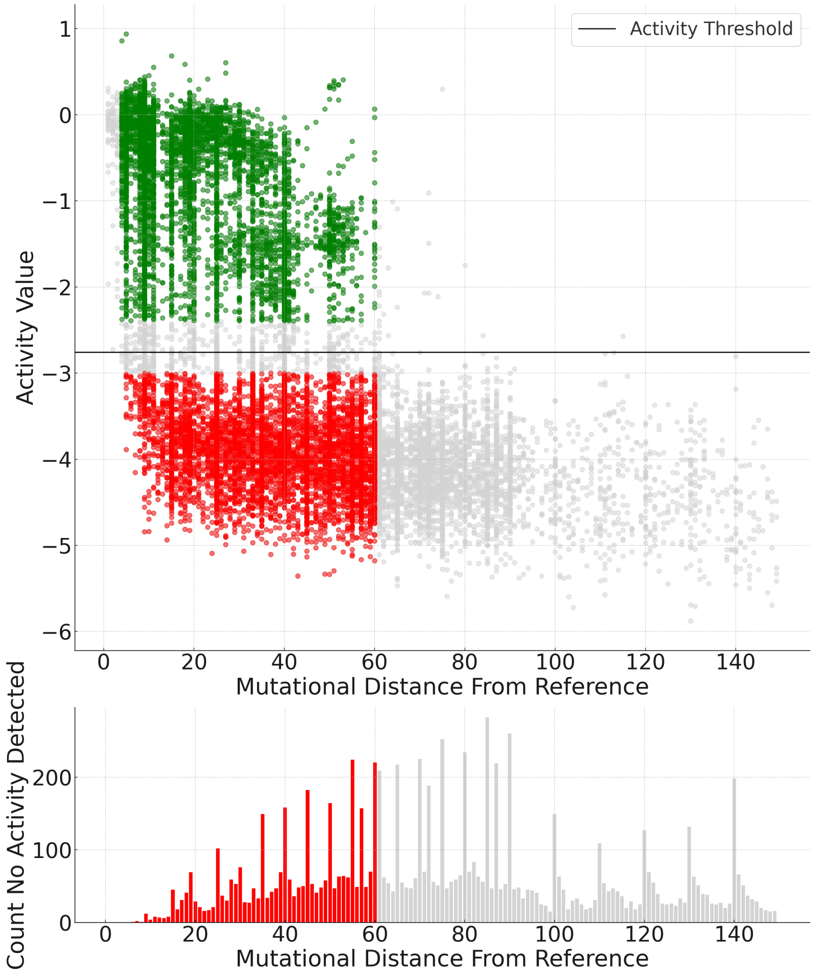


FIG. S3: *Self-splicing experimentally annotated dataset from [6] The top plot shows the activity versus distance from the reference sequence for this dataset. The horizontal line represents the activity threshold (-2.76) set in [6], where sequences with activity above the threshold are considered active, while those below are considered inactive. Colored sequences (green/red) represent the sequences used in the $\mathcal{D}_T$ reintegration dataset. Green dots represent reintegrated positive examples ($\mathcal{D}_T^+$), and red dots represent reintegrated negative examples ($\mathcal{D}_T^-$). Sequences with activity values close to the threshold ($-3.00 < x < -2.40$) were excluded from reintegration to avoid ambiguous signals. The bottom plot shows the number of sequences with an activity of $-\infty$ for each bin of distance.*

#### S4.2. Training Procedure and choice of $\lambda$

For the training of the two reintegrated models on the RNA group I intron self-splicing ribozymes, we employed a previous version of the adabmDCA algorithm, implemented in JAX. To ensure convergence, we set any negative

effective frequencies to zero, consistent with our approach in all previous cases. The training procedure followed the standard Pytorch adabmDCA protocol, utilizing 10,000 Monte Carlo chains and 10 sweeps per gradient update, but with a reduced learning rate of 0.01. Training was terminated once the Pearson correlation between the empirical and training correlation matrices, $C_{ij}^{\text{empirical}}(a,b)$ and $C_{ij}^{\text{training}}(a,b)$, reached a threshold of 0.95. It is important to note that the non-reintegrated original DCA model, was trained using the eaDCA algorithm. However, in this case, the eaDCA procedure failed to converge within $10^4$ training steps, necessitating the use of the adabmDCA approach.
The choice of $\lambda$ for the two reintegrated models is not straightforward for two main reasons. First, we are measuring actual experimental activity, which means we cannot run multiple experiments to tune the parameter. Second, we used a slightly more sophisticated adjustment function.
For the REINT case, $\lambda$ was set to 5000. The number of sequences in $\mathcal{D}_T^+$ is 5455, and the number of sequences in $\mathcal{D}_T^-$ is 8644. Accordingly, the weight assigned to positively reintegrated sequences is $w = 1/5455$, while for negatively reintegrated sequences, it is $w = 1/8644$. The value of $\lambda = 5000$ was chosen such that $w \cdot \lambda \approx 1$, a setting that consistently produced good results in computational examples.
In the case of the REINT BS0 model, the average size of the 14 $\mathcal{D}_T^+$ bins was 390 sequences, while the 14 $\mathcal{D}_T^-$ bins averaged 620 sequences. Ideally, a $\lambda$ value of around 400–500 should have been selected for the same reason as before, however, convergence issues arose during training. To address this, we selected the largest $\lambda$ value that led to convergence within 3 hours (with a learning rate of 0.01), resulting in $\lambda = 100$.

#### S4.3. Experimental validation of activity of predicted RNA sequences

The goal of the experimental procedure was to identify from the designed variants the active one. The selection procedure was based on the self-splicing-like assay (Fig. S4A) [6]. Initially, exon sequence A is covalently attached to the 3'-end of the ribozyme molecule. During the first step of reaction, ribozyme binds to "substrate S1" and through recombination events forms the covalent link between "substrate S1" and A RNA fragments. Then the substrate detaches the formed "complex S1-A" and binds to the new "substrate B-S2". After, the ribozyme covalently attaches the "S2" part of the substrate to the 3'-end of itself. The variants that have attached the "S2" part can be specifically selected during the library preparation step. The frequency of each RNA variant in the initial pool was used as the reference value and taken into account in the final calculation of the activity of each variant (Fig. S4B). This method was successfully applied to evaluate the activity of designed variants of the *Azoarcus* ribozyme [6].

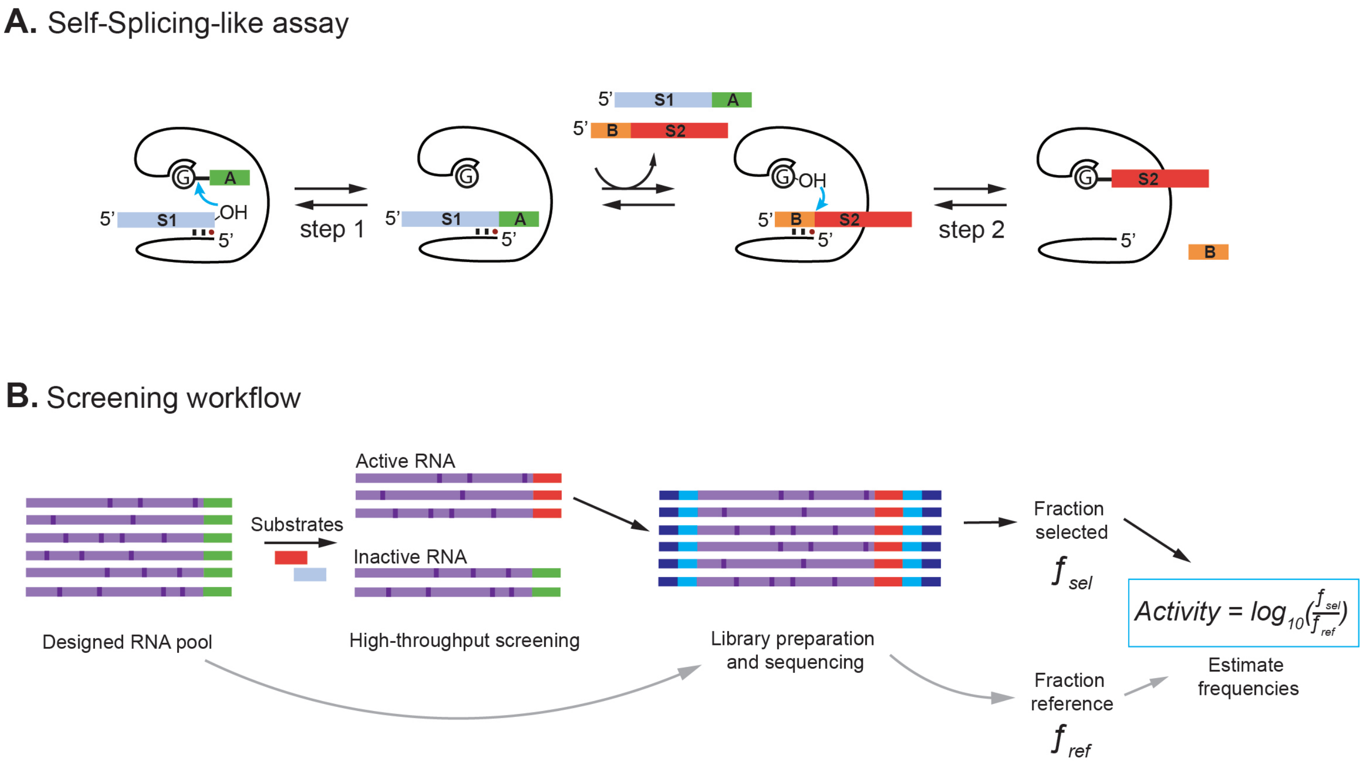


FIG. S4: *High-throughput screening of functional RNA. (A) Self-splicing-like assay. (B) Scheme of the high-throughput screening of functional RNA sequence. Adapted from [6].*

#### 1. *RNA production*

Designed RNA sequences were ordered as corresponding ssDNA templates with the exon sequences at the 3'-end ('AATCCGTTGGTGCTG'), and the T7 promoter at the 5'-end ('TAATACGACTCACTATA'). The ssDNA pool contained 12000 sequences was purchased from Twist Bioscience. The primers and RNA substrates were ordered from the Integrated DNA Technologies (IDT).

All reactions were performed using RNA DNAse/RNAse-Free Water (UltraPureTM Distilled Water, Invitrogen). The DNA pool was amplified by PCR (16 cycles) using the KAPA HiFi HotStart ReadyMix (Roche) and primers F_PCR and R_PCR, after which the samples were purified using the NucleoSpin Gel and PCR Cleanup kit (Macherey-Nagel). The RNA pool was transcribed from the amplified DNA pool using the HiScribe T7 High Yield RNA Synthesis Kit (New England Biolabs, NEB) for 4h at 37°C in a dry bath (MyBlockTM mini dry bath). Afterwards, DNAse I treatment was performed with 10U of DNAse I in 1X DNAse I Buffer (NEB), samples were incubated for 10min at 37°C in a dry bath.

An equal amount of phenol-chloroform (Ambion) was added to the reaction. Samples were vortexed for 1min and centrifuged for 4min at 11000rpm in a MiniSpin centrifuge (Eppendorf). The upper phase was transferred in 0.1 volume of 3M Sodium acetate (Sigma) with subsequent addition of 2.5 volumes of cold 100%. RNA samples were precipitated overnight at -20°C.

The sample was centrifuged for 1h at 14rpm at 4°C (Centrifuge 5418 R, Eppendorf). After, all supernatant was removed and the pellet was gently washed with 150$\mu$l of 70% cold ethanol twice. The left ethanol was evaporated using a vacuum concentrator (Concentrator Plus, Eppendorf). The dry pellets were resuspended in 40$\mu$l of water. After 50$\mu$l of loading dye (90% formamide, 100 mM EDTA (éthylènediaminetétraacétique), 0.1% xylene cyanol, 0.1% bromophenol blue) was added to each sample.

The polyacrylamide gel (20cm $\times$ 20cm) with 8M urea was prepared using the ROTIPHORESE DNA sequencing system (Carl Roth). The samples were loaded onto the 8% urea PAGE. The gel was run for 1h at 420V. The gel covered by transparent plastic film was placed on Thin Layer Chromatography sheets topped with silica gel (Macherey-Nagel) and illuminated by a UV lamp at 254nm. The sections of the gel corresponding to the produced RNA were then cut with a sterile scalpel and transferred to a new 1.5ml Eppendorf tube. The gel pieces were crushed using a

1ml pipette tip. To each sample 500 $\mu$l of 0.3M sodium acetate was added. The tubes were incubated at 26°C for 5h at 450rpm in a ThermoMixer Dry Block (ThermoMixer F1.5, Eppendorf). After incubation, the upper phase was transferred to columns with a 0.22$\mu$m filter (Corstar) and centrifuged for 4 minutes at 11,000rpm. Then, 2.5 volumes of cold 100% ethanol were added to each solution. Samples were left at -20°C overnight.
The sample was centrifuged for 1h at 14 rpm at 4°C. The supernatant was removed and the pellet was washed two times with 150 $\mu$l of cold 70%. The residual ethanol was evaporated using a vacuum concentrator. The dry pellets were resuspended in 20$\mu$l of water. The final concentration was measured with the spectrophotometer NanoDrop One (Thermo Scientific).

#### 2. *Self-splicing-like assay*

The self-splicing-like reaction was perfomed according to the following protocol, 1$\mu$M of RNA pool were incubated with 25$\mu$M of "substrates S1" and "B-S2" in a reaction buffer (30mM EPPS pH7.5, 60mM $MgCl_2$) at 37°C for 1h in a final volume of 14$\mu$l. The reaction was quenched by adding EDTA to final concentration of 60mM, and cleaned using the Monarch RNA cleanup kit (New England Biolabs) with an adjusted volume of 100%ethanol (75$\mu$l ) and binding buffer (75$\mu$l). The sample were eluted in 12$\mu$l of water.
A control experiment without substrate addition was conducted to correct for biases in the relative quantity of each synthesized ribozyme within the corresponding pool. The initial RNA pool was diluted to 1$\mu$M in a reaction buffer with a final volume of 14$\mu$l. Without incubation, the reaction was directly quenched by adding EDTA to a final concentration of 60mM. Samples were purified using the Monarch RNA Clean up Kit following the same protocol as for the self-splicing-like assay.

| Name | Sequence | Type |
|---|---|---|
| S1 | rCrGrCrGrArArUrUrArArCrGrCrGrArCrArArCrArU | RNA |
| B-S2 | rGrGrCrArUrArArCrUrUrCrArArArUrArUrCrUrUrCrGrGrArArCrUrCrA | RNA |
| F_PCR | TAATACGACTCACTATAGTG | DNA |
| R_PCR | CAGCACCAACGGATTCC | DNA |
| RT_S2 | TGAGTTCCGAAGATATTTGAAGTTCC | DNA |

TABLE S5: *List of RNA and DNA oligos*

#### 3. *Library preparation and sequencing*

The RNA samples were prepared for sequencing using NEBNext Ultra II Directional RNA Library Prep Kit for Illumina (NEB). At the first reverse transcription step, the primer RT_S2, which is complementary to the S2 part of the substrate, was added to the RNA sample that had undergone a self-splicing-like reaction. Similarly, for the control reaction, the primer R_PCR, which is complementary to the exon part A, was used. During PCR amplification step each sample was barcoded using NEBNext Multiplex Oligos for Illumina (Dual Index Primers Set 1, NEB).
The final library sequenced on a NovaSeq SP flow cell (2x250 nts, 2x800 M reads) in paired ends and with 20% of PhiX by the NGS platform at Institut du Cerveau et de la Moelle épinière (ICM, Paris) or Institut Curie (Paris).

#### 4. *Experimental Activity Scores from Sequencing Data*

To estimate the experimental activities from sequencing data, we followed the exact same procedure as in [6]. We computed the frequencies of designed sequences under two conditions:

1. the reference condition, prior to the catalytic reaction,

2. the reacted condition, where the substrate was mixed with the designs and incubated.

For both conditions, we mapped each paired-end read to the closest designed sequence using **BLAST** (version 2.12) [7]. Reads were retained only if they covered at least 70% of the mapped designed sequence with full identity.
We first computed the frequencies $f_{\mathrm{ref}}$ of designs before the catalysis, which allowed us to quantify the bias in the initial synthesized pool of RNA molecules. Next, we computed the frequencies $f_{\mathrm{sel}}$ of designs in the reacted sample.

To determine $f_{\text{sel}}$, we counted reads where the substrate was attached immediately after the 3' end of the design, indicating successful excision of the exon and subsequent ligation of the substrate.
The experimental activity was then calculated as:

$$\text{act} = \log_{10}(f_{\text{sel}}/f_{\text{ref}})$$

Analysis of reverse reads was sufficient for computing the activity score. Designs with fewer than 5 reads in the pre-catalysis pool were excluded from the analysis to avoid ambiguity. Sequences with $f_{\text{ref}} > 0$ and $f_{\text{sel}} = 0$ were classified as inactive.

#### 5. *Comparability with Previous Experiments*

Since we compare the results of our experiment against those of previous experiments [6], even if the experimental assay is identical, it is essential to address the issue of comparability of the results. Fortunately, the experimental assay presented is highly reproducible and the results of different experimental pools can be easily compared.
The experimental data used for our reintegration in [6] is already the outcome of three independent assay pools. In [6], they used 355 overlapping sequences tested in these pools to align the resulting experimental values. The measured experimental activity $\log_{10}(f_{\text{sel}}/f_{\text{ref}})$ exhibits a correlation of up to 99% across different pools, and aligning the results only requires the introdcution of an additive constant. Specifically, if $\text{act}_i$ represents the activity measured in the $i$-th pool, then:

$$\text{act}_i = \log_{10}(f_{\text{sel}}/f_{\text{ref}}) + \alpha_i,$$

where $\alpha_i$ is the constant used to align the values.
In our case, since alignment with the $P^1$ sequences is crucial, we used the same 355 overlapping sequences to align the activity values with those from the $P^1$ pool in [6]. The results of this alignment are shown in Figure 4. The comparison reveals an almost perfect match, with a linear regression slope $m = 1.02$ and intercept $q = 0$ ($R^2 = 0.96$). Thus, the chosen $\alpha_i$ was set to the same value as the $P^1$ pool from [6] ($\alpha_i = -0.5103$), ensuring that the activity values are directly comparable to those presented in [6].

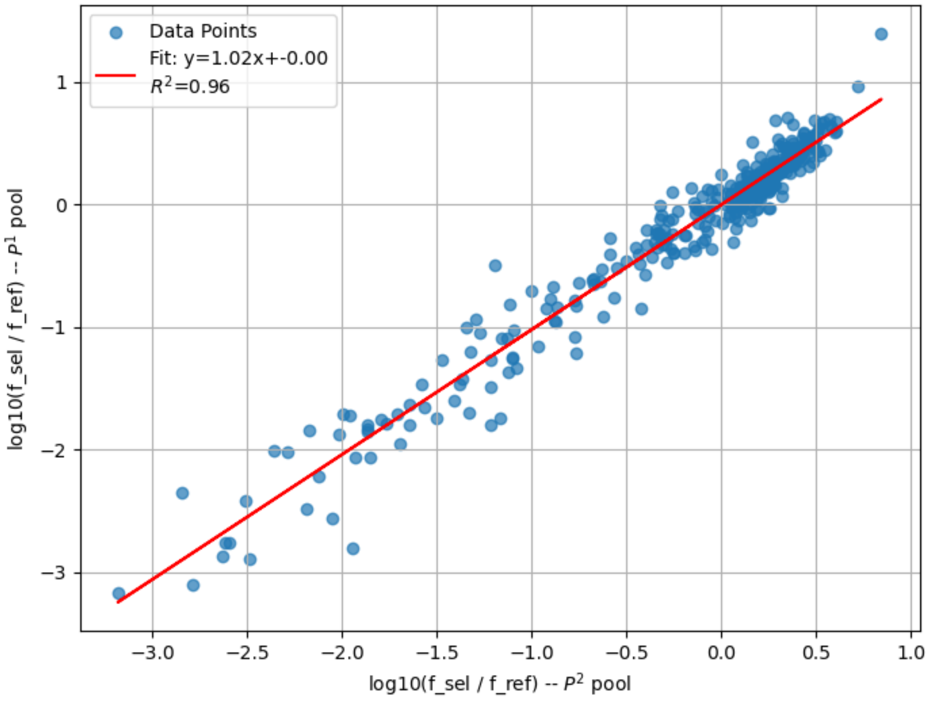


FIG. S5: *Comparison of* $\log_{10}(f_{sel}/f_{ref})$ *between the* $P^1$ *pool [6] (y-axis) and* $P^2$ *pool (x-axis). The linear regression analysis reveals a near-perfect match with a slope of* 1.02*, an intercept of* 0*, with an* $R^2$ *value of* 0.96.

### 6. Group I Intron: Tables and Violin Plot of key metrics

Table S6 summarizes key metrics, including active fractions of sequences at different distances. To better visualize these metrics, violin plots in Figures S6 and S7 show, respectively, the distribution of intra-dataset distances and minimum distances to the reintegrated dataset.

| Model (Distance) | Active % | $D_{P^2-P^2}$ | $D_{P^2-\mathcal{D}_T^+}$ |
|---|---|---|---|
| DCA $P^1$ (30) | 43.3 | 35.5 | 20.8 |
| REINT BS0 (30) | 99.0 | 6.4 | 5.6 |
| DCA $P^1$ (45) | 6.7 | 51.3 | 31.8 |
| REINT (45) | 63.7 | 8.7 | 6.5 |
| REINT BS0 (45) | 52.0 | 10.0 | 7.6 |
| DCA $P^1$ (55) | 2.0 | 59.4 | 38.3 |
| REINT (55) | 14.4 | 11.0 | 14.2 |
| DCA $P^1$ (60) | 2.0 | 62.8 | 43.1 |
| REINT (60) | 3.3 | 12.5 | 18.6 |
| REINT BS0 (60) | 23.6 | 15.1 | 13.8 |
| DCA $P^1$ (65) | 0.0 | 67.7 | 47.2 |
| REINT (65) | 2.8 | 14.2 | 23.1 |
| DCA $P^1$ (70) | 0.0 | 70.4 | 50.3 |
| REINT (70) | 0.6 | 18.1 | 28.0 |
| DCA $P^1$ (75) | 0.0 | 75.5 | 55.7 |
| REINT BS0 (75) | 0.7 | 33.3 | 25.1 |

TABLE S6: *Active fraction, average intra-dataset distance ($D_{P^2-P^2}$), and average minimum distance from the positively reintegrated dataset ($D_{P^2-\mathcal{D}_T^+}$) for mutational distances 30, 45, 55, 60, 65, 70, and 75 from the reference sequence.*

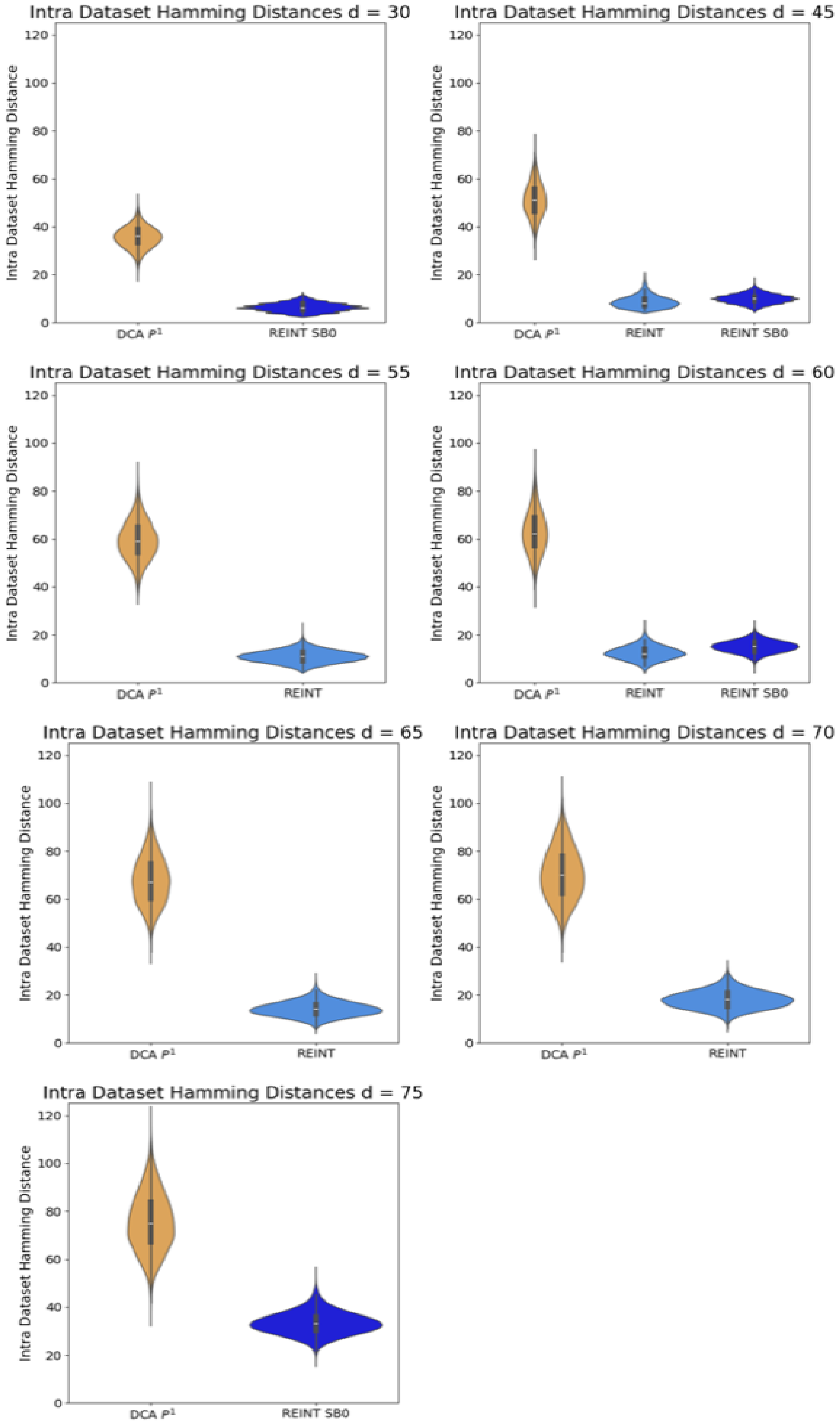


FIG. S6: *Violin Plot of intra-dataset distance ($D_{P^2\_P^2}$) for mutational distances 30, 45, 55, 60, 65, 70, and 75 from the reference sequence.*

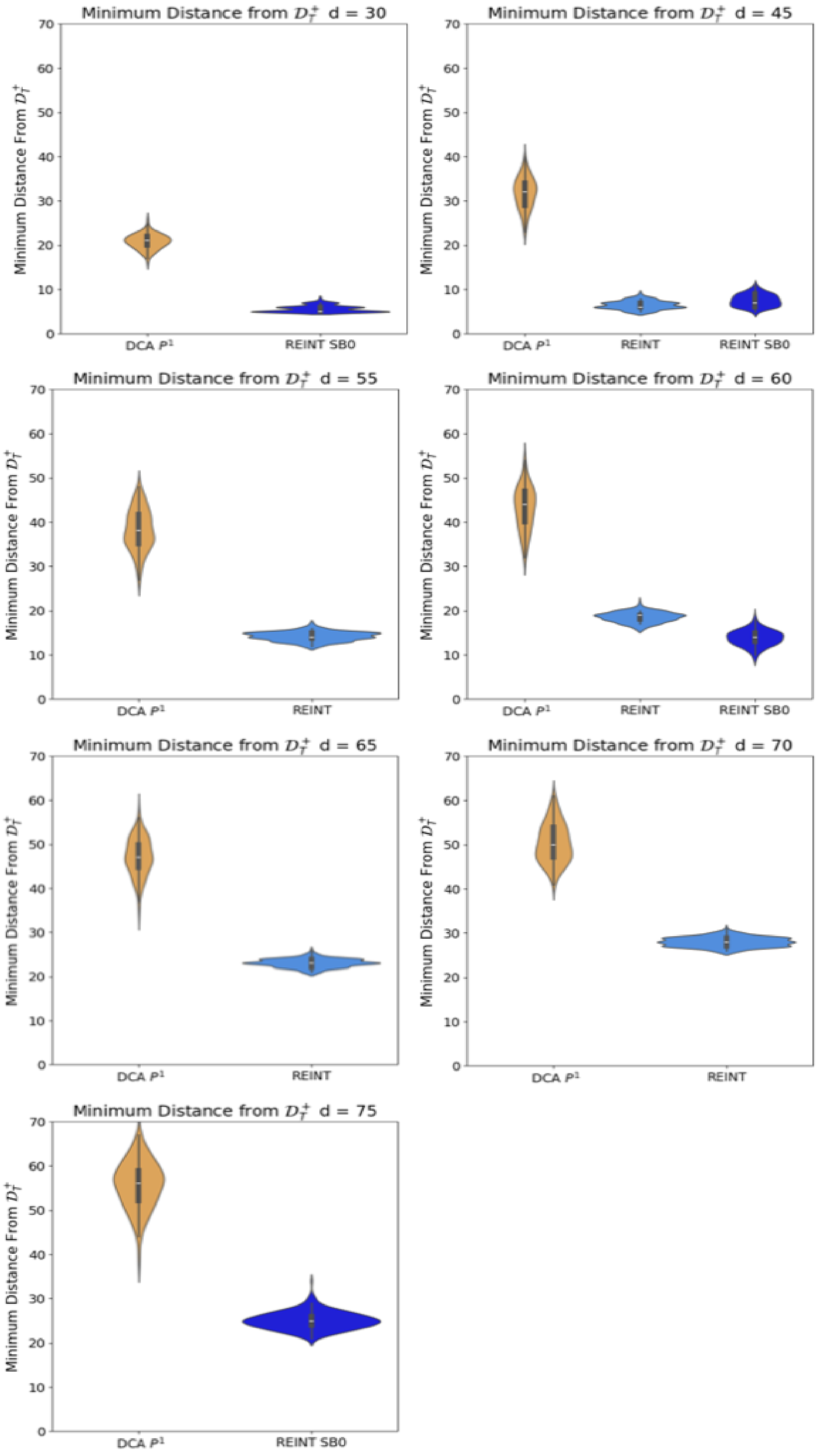


FIG. S7: *Violin Plot of minimum distance from the positively reintegrated dataset ($D_{P^2-\mathcal{D}_T^+}$) for mutational distances 30, 45, 55, 60, 65, 70, and 75 from the reference sequence.*

[6] C. N. Lambert, V. Opuu, F. Calvanese, F. Zamponi, E. Hayden, M. Weigt, M. Smerlak, and P. Nghe, "Expanding the space of self-reproducing ribozymes using probabilistic generative models." July 2024.

[7] S. F. Altschul, W. Gish, W. Miller, E. W. Myers, and D. J. Lipman, "Basic local alignment search tool," *Journal of Molecular Biology*, vol. 215, pp. 403–410, Oct. 1990.
